# CA2/3-dependent stability of frontoparietal mnemonic representations predicts episodic deficits in human amnesia

**DOI:** 10.1101/2025.02.19.638996

**Authors:** Thomas D. Miller, Alice L. Hickling, Yan I. Wu, Joseph H. Zhou, Adam E Handel, Ester Coutinho, Thomas A. Pollak, Michael S. Zandi, Clive R. Rosenthal, Eleanor A. Maguire

**Affiliations:** Functional Imaging Laboratory, University College London, London, UK; UCL Queen Square Institute of Neurology, National Hospital for Neurology and Neurosurgery, Queen Square, London, UK; UCL Medical School, University College London, London, UK; Oxford Autoimmune Neurology Group, Nuffield Department of Clinical Neurosciences, University of Oxford, Oxford, UK; Department of Neurology, John Radcliffe Hospital, Oxford University Hospitals NHS Foundation Trust, Oxford, UK; Center for Neuroscience and Cell Biology, Universidade de Coimbra (CNC-UC); Centre for Innovative Biomedicine and Biotechnology (CiBB), Universidade de Coimbra; Católica Medical School, Universidade Católica Portuguesa; Department of Psychosis Studies, Institute of Psychiatry, Psychology and Neuroscience, King’s College London, London, UK; Nuffield Department of Clinical Neurosciences, University of Oxford, Oxford, UK

**Author notes:** contributed equally to this work (Clive R. Rosenthal, Eleanor A. Maguire).

## Abstract

The hippocampus reconstructs past experiences by integrating sensory, perceptual, and conceptual information across a cortico-hippocampal autobiographical memory network (AMN). Here, in 18 human participants with amnesia, we decoded the effects of bilateral focal hippocampal damage on distinct autobiographical representations using representational dissimilarity matrices (RDMs). Hippocampal pathology resulted in: (1) impaired generalised episodic memory retrieval RDM model fit in the left angular gyrus; and, (2) reduced distinct episodic memory RDM model fit and trial-by-trial voxel-representational stability in the right angular gyrus, right orbitofrontal cortex, and right inferior frontal gyrus. RDM model fits and mnemonic stability were predicted by total CA23 volumes. Trial-by-trial retrieval stability within the right orbitofrontal cortex and right inferior frontal gyrus predicted autobiographical memory performance, providing the first direct neural correlate between hippocampal dysfunction, altered mnemonic representations, and amnesia. The results demonstrate that hippocampal-mediated amnesia results from impaired mnemonic representational distinctiveness and stability within the AMN.

---

Autobiographical recollection, a fundamental component of episodic memory, involves the reconstruction of past experiences by integrating sensory, perceptual, and conceptual information into cohesive personal narratives.^1–5^ This integration relies on a large-scale network involving parietal regions, such as the angular gyrus and dorsal/ventral parietal cortex, which enable multimodal sensory processing and contextual framing within the autobiographical memory network (AMN).^5–13^ Temporal-frontal interactions, involving the hippocampus and ventromedial and medial prefrontal cortex, facilitate the integration of conceptual elements of recollection, including schema formation, self-processing, and perceptual abstraction.^14–20^

The hippocampus is also essential for forming coherent mental representations required for episodic memory, integrating both unimodal and multimodal inputs from broader processing networks.^1–5,21^ Lesion studies underscore the essential role of the hippocampus in retrieving episodic details, as hippocampal damage often leads to episodic, and especially autobiographical, amnesia.^22–27^ Functional MRI (fMRI) shows that hippocampal damage, and wider medial temporal lobe (MTL) pathology, disrupts the dynamic interactions between the hippocampus and other AMN regions, which impairs connectivity and memory processes needed for autobiographical retrieval.^28–31^

Although prior fMRI human hippocampal lesion studies have been informative, their interpretation is often complicated by the inclusion of individuals with anatomical damage that extends beyond the hippocampus to other medial temporal lobe structures and associated network nodes.^32–34^ Focal models of hippocampal damage in humans are rare,^22,31,35–37^ limiting group-level behavioural and fMRI assessments of its impact on autobiographical memory. An additional concern is that prior fMRI studies of participants with hippocampal damage used univariate analyses, thereby failing to capture important details about the content and structure of reinstated memories. In contrast, more recent work using multivariate analyses in neurologically-intact humans suggests that the hippocampus coordinates the reinstatement of distributed neural patterns associated with an event during its initial encoding period.^38–41^ This highlights the need for more sophisticated analyses^12,42–44^ in studies of hippocampal amnesia.

Here we investigated the effects of bilateral focal hippocampal damage, associated with episodic amnesia, across the AMN, decoding how this damage altered the representational content of pre-morbid, recently acquired autobiographical memories during retrieval. Eighteen human participants with amnesia, secondary to a single aetiology, leucine-rich glioma-inactivated-1-limbic encephalitis (LGI1-LE),^37,45–47^ were recruited and compared with a group of 18 matched control participants. Anatomical MRI-based characterisation confirmed that the damage was bilateral and confined to the hippocampus in all of the participants with amnesia (bilateral volume loss was confined to CA1 and CA23 subfields, mean reductions = -18% and -41%, respectively). These novel results of focal hippocampal damage align with previous findings in the same aetiology.^31,37,48–55^

We used then fMRI to compare autobiographical memory retrieval in the participants with hippocampal damage-mediated amnesia with the control group. The retrieval protocol was optimised for decoding representational content. Episodic and semantic details associated with each autobiographical memory retrieved in the scanner were interrogated using an objective, parametric, quantitative method, the Autobiographical Interview (AI) procedure.^56^ The AI was chosen because it has demonstrated sensitivity to detecting and characterising episodic amnesia linked with hippocampal pathology.^24,31,37,57–59^ Consistent with prior results,^31,37^ we found that autobiographical memories in the hippocampal amnesia group were associated with a loss of internal (episodic) details of autobiographical memories, but not external (semantic) details. Accordingly, we use the term ‘episodic’, rather than ‘autobiographical’ memory, because this better reflects the re-experiential component of autobiographical memories that are specifically impaired after bilateral hippocampal damage.

Unlike previous studies of hippocampal damage that focused on overall univariate activity changes, we measured the consistency (stability) of the neural pattern activity each time a memory was retrieved. Moreover, to decode specific memories, we calculated representational dissimilarity matrices (RDMs) based on memory-specific neural response patterns. RDMs (an approach within multivoxel pattern analysis (MVPA) have been used to characterise voxel-level signal patterns across multiple voxels in response to specific memories, revealing fine-grained representational structures and enabling discrimination between individual memories,^42,60^ even when the overall activity levels in a region remains relatively constant.^61,62^ Application of MPVA based pipelines in healthy adults have linked hippocampal activity with the successful differentiation, reinstatement, and retrieval of specific episodic memories.^12,43,44,63–68^ In the current study, RDMs offered two advantages over other frameworks within MVPA:^69^ (1) we obtained quantitative measures of dissimilarities to compare the neural data with different experimental hypotheses about the type of information being represented in a brain region during mnemonic retrieval;^14,42,61,70,71^ and, (2) we were able to quantify the consistency of voxel activity patterns across multiple stimulus presentations, yielding a measure that indicated the stability of specific personal event memories.^14,69–71^

Hippocampal damage is also known to impair the differentiation of memories,^72–76^ which aligns with findings from healthy adults that has identified the hippocampus, especially the dentate gyrus and CA3 subfields, with unique and discriminable activity patterns for similar stimuli (such as memories).^77^ Together, these observations lead to the hypothesis that a loss of pattern separation, and hence mnemonic discriminability, may partially explain episodic memory deficits following hippocampal damage.

To investigate how hippocampal damage affects the differentiation of episodic memories, we tested whether RDMs would show reduced discriminability (or distinctiveness)^69^ between episodic memories in the group with hippocampal amnesia compared to healthy controls. Specifically, we predicted decreased dissimilarity between neural representations of these episodic memories for the group with hippocampal amnesia. Our second hypothesis was that these effects would be most pronounced in AMN regions that are closely connected to the hippocampus, given that lesions involving network hubs like the hippocampus can disrupt functional connectivity of local and non-local brain regions.^31,78–81^ Our third hypothesis was that RDM-derived neural measures in affected AMN nodes would predict the amount of episodic detail remembered on the basis that differentiated neural representations enhance richly-detailed remembering and reduce interference from similar memories.^38,82,83^ This aligns with evidence that RDM-derived neural measures can serve as a neural code informing behavioural responses.^31,37,64,84–86^ Our fourth hypothesis was that the extent of hippocampal pathology would be linked to representational stability, with greater damage leading to less stable representations, evidenced by higher dissimilarity across trials for the same memory and/or more similarity across different memories.^69^ Such instability in the group with hippocampal amnesia would suggest a less consistent representational structure, signalling degraded memory retrieval.^69^ Our study therefore sought to provide new insight into how hippocampal dysfunction is expressed in the neural systems that support autobiographical memory.

**Figure 1.**
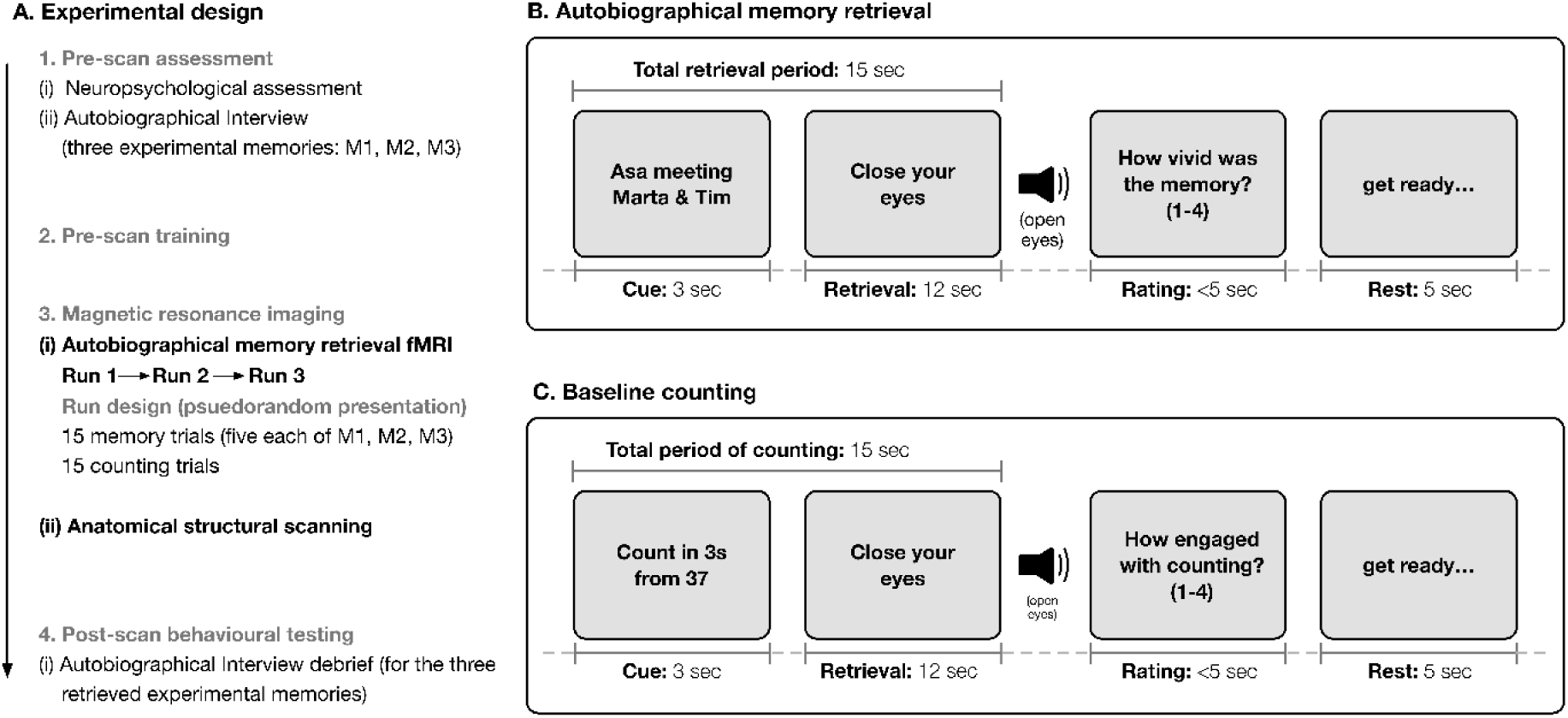
Experimental design. Episodic memory decoding task protocol. (Panel A) The experiment was comprised of four separate phases (1). Two-weeks prior to the scan date, all participants underwent extensive neuropsychological assessment and the Autobiographical Interview (AI) procedure to acquire three personal episodic memories that would be retrieved during the experimental scanning. (2-3) On the day of scanning, all participants underwent extensive (2) pre-scan training and (3) intra-scan training (see Methods). Once in the MRI scanner, all participants undertook three fMRI runs comprised of two conditions (autobiographical memory retrieval and the baseline counting task). (4) Post-scan behavioural testing. Immediately after the end of two anatomical scans, a final debriefing on the experimental memories was conducted. Participants were asked to recount what they retrieved for each memory during the retrieval periods using the AI procedure. These debriefing interviews were recorded in a digital audio format and then transcribed for offline scoring. fMRI trial types (Panel B) Autobiographical memory retrieval. Participants were presented with a three-second verbal cue for a specific autobiographical memory. After the offset of the cue, the participants were prompted to then close their eyes and retrieve the memory with as much visual and verbal episodic detail as possible. The prompt to “close your eyes” remained on the screen for 12 seconds), while the participants continued to retrieve the cued memory. An auditory instruction to “Open your eyes” cued the end of the retrieval period. (Panel C) Counting trials. Participants were presented a starting number for three-seconds and then a specific interval to count up in 3s, followed by the same ‘Close your eyes’ screen prompt for 12 seconds. After each memory retrieval trial and counting trial, the participants were then asked to rate either ‘how vivid was the recollection?’ or ‘how engaged were you with the task?’ (see Methods), respectively, and indicate their response using an MRI-compatible button box. All trials were followed by a five-second rest period before the next trial. No inter-trial jitter was used given the length of retrieval trials. Immediately after the completion of the fMRI experimental phase, each participant underwent two anatomical scans.

## RESULTS

First, we report the results of structural brain imaging analyses to determine whether the damage was limited to the hippocampus and which hippocampal subfields were damaged in the group with hippocampal amnesia. Second, we report episodic and semantic memory retrieval performance in the group with hippocampal amnesia, using the degree of hippocampal subfield loss to predict the severity of retrograde episodic amnesia.^31,37^ Third, we report the fMRI RDM results to determine whether neural response model fit in the AMN differed in the amnesic and control groups for measures of (i) general episodic memory and (ii) specific episodic memories. Fourth, we then report the stability of specific memories using the averaged dissimilarity scores from the first-level RDM analysis (calculated as 1 - Fisher-transformed Pearson correlation scores), and how these stability measures related to the degree of hippocampal subfield damage and episodic memory performance.

### Selective bilateral hippocampal subfield volume loss

The group with hippocampal amnesia had focal hippocampal subfield pathology in the CA23 and CA1 subfields, according to quantitative three-dimensional whole-hippocampal volumetry of six hippocampal subfields (see Figure 2 and Supplementary Table 4). A three-way 2 (group: amnesic, control) x 2 (side: left, right) x 6 (subfield: dentate gyrus, CA23, CA1, subiculum, pre/parasubiculum, uncus) mixed-model ANOVA [with Mauchly’s test demonstrating that the assumption of sphericity had been violated for subfield (χ^2^_(14)_ = 59.583, *p*<0.001) and subfield by side (χ^2^_(14)_ = 78.452, *p<* 0.001), therefore, degrees of freedom were corrected using Huynh-Feldt estimates ε = 0.85 and 0.64, respectively] demonstrated there were significant main effects of group [*F*_(1,34)_ = 12.536, *p* = 0.001], side [*F*_(1,34)_ = 13.537, *p<*0.001], and subfield [*F*_(4.504,153.152)_ = 415.019, *p<*0.001]. A significant two-way interaction was observed between group and subfield [*F*_(4.504,153.152)_ = 4.784, *p*<0.001]. The three-way interaction was not significant [*F*_(3.679,125.082)_ = 0.894, *p* = 0.463]. Planned comparisons demonstrated that this between-group volume loss was isolated to the CA23 and CA1 subfields (*F*_(1,34)_ = 129.639, *p*<0.001, η^2^ = 0.792 and *F*_(1,34)_ = 15.330, *p*<0.001, η^2^ = 0.311, respectively; Bonferroni-corrected *p* = 0.0083; see Figure 3A&B).

**Figure 2.**
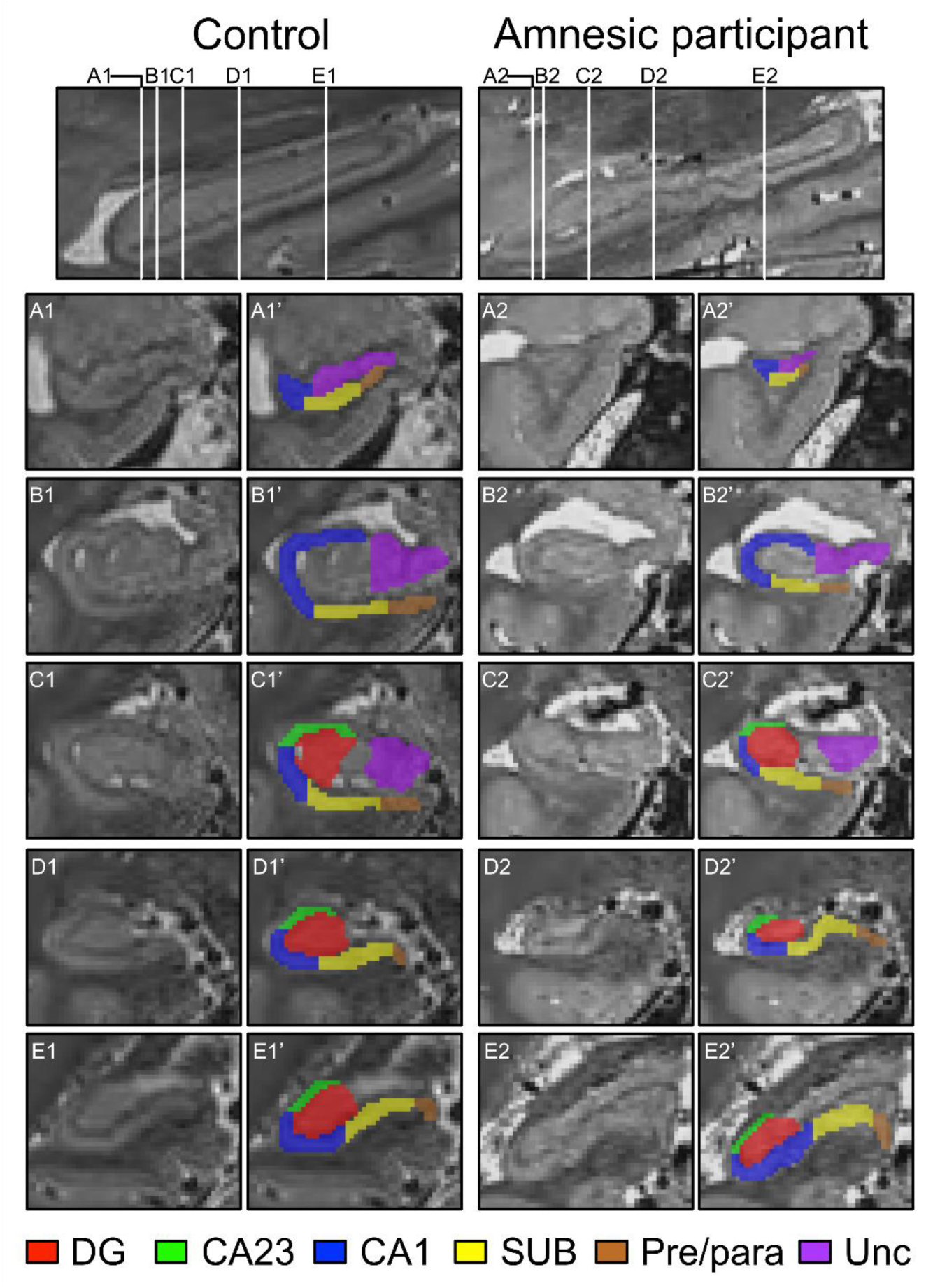
Quantitative 3.0-Tesla (0.5mm^3^ spatial resolution) 3D T2-weighted turbo spin echo images coregistered and averaged with Rician noise estimation and oracle-based discrete cosine transform were used to conduct whole hippocampal semi-automated volumetry of six hippocampal subfields (CA23, CA1, dentate gyrus, subiculum, pre/parasubiculum, and the uncus) in the hippocampal amnesia group and in a control group of participants. Denoised sagittal images on the first row illustrate the full longitudinal axis of hippocampi in a (matched) control participant and participant with amnesia. Each of the white lines (A-E) on the sagittal view of the hippocampus corresponds to six example coronal locations along the anterior-posterior axis. Coloured shading on the left hand side coronal images on rows A-E under the matched control and participant with amnesia provide examples of the results of applying the hippocampal segmentation protocol in the control and amnesic participant (denoted as,’). CA1: cornu Ammonis 1; CA23: cornu Ammonis 2 & 3; DG: dentate gyrus; Pre/para: pre/parasubiculum; SUB: subiculum; Unc: uncus.

**Figure 3.**
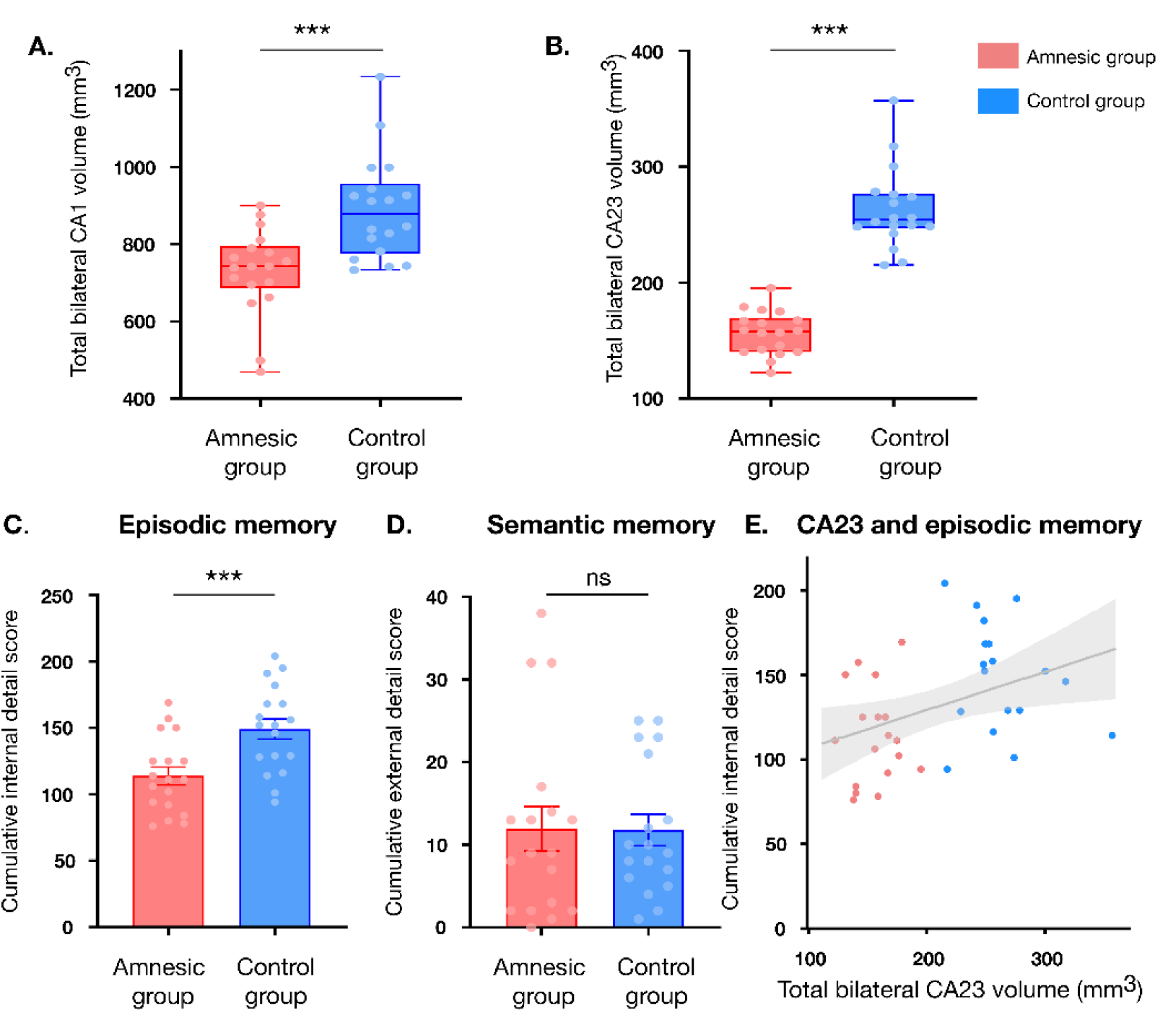
CA23 atrophy and impairment of cumulative episodic memory performance in the group with hippocampal amnesia (i.e., single LGI1-LE aetiology group). Hippocampal segmentation was based on 3D T2-weighted turbo spin-echo images coregistered and averaged with Rician noise estimation and oracle-based discrete cosine transform. Results from hippocampal subfield segmentation in the group with hippocampal amnesia (n = 18) relative to the control group. Results from the analyses of hippocampal segmentation are plotted in panels A-B. (A) total CA1 and (B) total CA23 volumes box-and-whiskers plots in both groups (representing the maximum and minimum values, with the horizontal line denoting the group mean). An omnibus ANOVA revealed a significant (alpha criterion Bonferroni-corrected *p* = 0.0083) reduction in-total CA1 [mean reduction = 18.13%; *F*_(1,34)_ = 15.330, *p*<0.001, η^2^ = 0.311] and CA23 [mean reduction = 40.53%; *F*_(1,34)_ = 129.639, *p*<0.001, η^2^ = 0.792] volumes in the group with hippocampal amnesia compared to the control group. Results from the analyses of the data obtained from the Autobiographical Interview (AI) procedure are plotted in panels C-D. Cumulative (summed across the three experimental memories) episodic memory performance detailing both (C) internal (episodic) and (D) external (semantic) details are each plotted as a function of group. The group with hippocampal amnesia exhibited a significant deficit in remembering internal (episodic) details relative to controls [*F*_(1,34)_ = 12.055, *p*<0.001, η^2^ = 0.262], whereas remembering external (semantic) details from the same episodic memories was comparable between the amnesic and control groups [*F*_(1,34)_ = 0.001, *p* = 0.973]. Error bars represent the standard error of the mean. (E) The scatter plot (with the best-fitting linear regression line) shows that total CA23 volume (independent variable) significantly predicts the total number of internal (episodic) details retrieved (dependent variable) across the three experimental memories [(*F*_(1,34)_ = 6.193, *p* = 0.018; *t* = 2.489, *p* = 0.018, R^2^ = 0.154, β_1_ = 0.393]. Total CA1 volume was not associated with this relationship (*t* = -0.238, *p* = 0.813). These results also replicate our previous findings in a new cohort of individuals with amnesia. Each dot represents a single data point, with red corresponding to a value for a participant in the group with hippocampal amnesia and blue corresponding to a value for a participant in the control group. *** denotes *p*<0.001. ns = not significant.

At the whole brain level, there was no significant loss or gain in normalised grey matter volume in the group with hippocampal amnesia, relative to controls, according to voxel-by-voxel contrast of normalized grey matter, conducted using a two-sample *t*-test thresholded at *p<*0.05 family-wise error corrected for multiple comparisons across the whole-brain. These results suggest that the memory deficits in the group with hippocampal amnesia are closely linked to specific hippocampal subfield damage, rather than widespread brain pathology.

### Subfield pathology is associated with retrograde episodic amnesia

In line with previous findings, neuropsychological assessment in the group with hippocampal amnesia demonstrated focal anterograde visual and verbal memory impairment when compared to the matched control population (see Supplementary data),^31,37,87^ alongside specific impairment in the retrieval of internal (episodic) but not external (semantic) memory details.^31,37^ An omnibus 2 (group: amnesic, control) x 3 (memory: experimental event memories) x 2 [memory detail type: internal (episodic), external (semantic)] mixed-model factorial ANOVA was conducted on units of information acquired from the AI for the experimental memories (Mauchly’s test confirmed that the assumption of sphericity had not been violated). Results revealed a significant main effects of group (*F*_(1,34)_ = 10.983, *p* = 0.002) and of memory detail type (*F*_(1,34)_ = 494.735, *p*<0.001) alongside a significant two-way interaction between group and detail type (*F*_(1,34)_ = 10.832, *p* = 0.002, η^2^ = 0.242), with planned comparisons (alpha criterion corrected for multiple comparisons, *p* = 0.025) revealing a significant loss of internal (episodic) memory detail in the amnesic group compared to the control group (*F*_(1,34)_ = 12.055, *p*<0.001, η^2^ = 0.262), whereas external (semantic) details were not significantly different (*F*_(1,34)_ = 0.001, *p* = 0.973; see Figure 3C&D).

### CA23 total volume loss predicted episodic amnesia

Next, we found that variability in total CA23 but not CA1 volume predicted internal (episodic) detail scores. This was revealed in a stepwise linear regression model with total CA23 and CA1 volumes entered as continuous independent variables and internal detail scores as the dependent variable. The model was statistically significant (*F*_(1,34)_ = 6.193, *p* = 0.018). Total CA23 volume selectively predicted the amount of internal (episodic) details retrieved (*t* = 2.489, *p* = 0.018, R²=0.154, β₁=0.393; see Figure 3E), whereas total CA1 volume was excluded from the model, even when the model was reversed (*t* = -0.238, *p* = 0.813).

### Hippocampal subfield damage selectively disrupts neural responses in the autobiographical memory network (AMN)

During the fMRI study, no between-group differences were observed for the intra-scan ratings (see Supplementary data), but the post-scan debrief test (see Methods) demonstrated a group difference in the retrieval of internal (episodic) details, replicating the pre-scanning behavioural results (see Supplementary data).

The fMRI data were first analysed using RDM analysis to test two hypothesised models of neural activity patterns during memory retrieval: (1) a model reflecting similarity across all episodic memory retrieval trials, independent of the specific episodic memory being remembered; and (2) a model depicting similarity across retrieval trials of the same specific episodic memory and dissimilarity between retrieval trials of different memories (see Figure 4).

**Figure 4.**
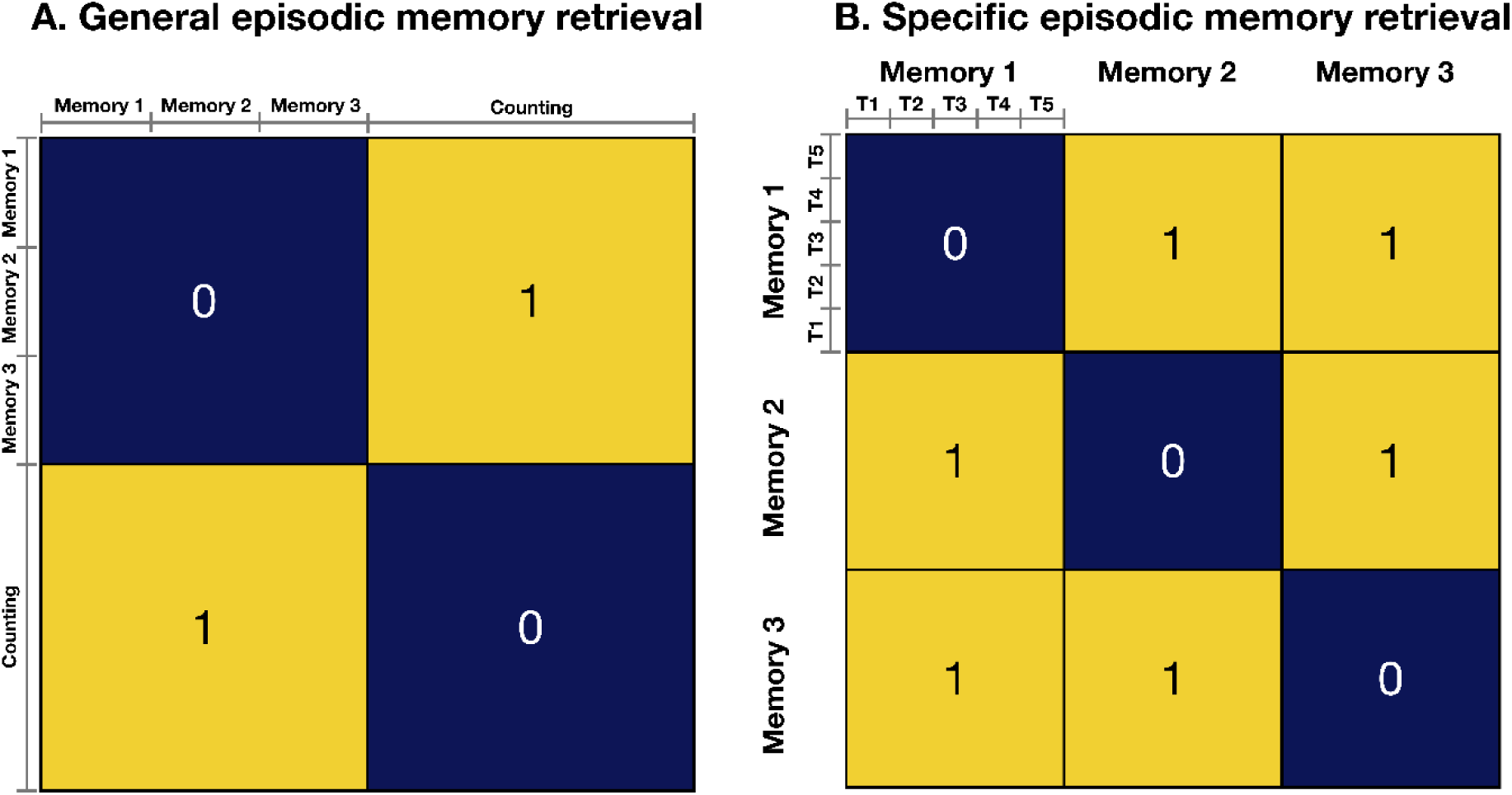
Model representational dissimilarity matrices (RDMs) based on hypothesised representational profiles. We performed a model fit analysis by comparing two RDMs within the representational patterns observed in the individual participants RDMs. Each specific memory in (A) and (B) is further subdivided into five trials per run (as shown in B), and there were also 15 counting trials per run. Final values were averaged across runs. These matrices were designed to test two key hypotheses: (A) whether focal hippocampal subfield damage resulted in between-group differences in representational content during episodic memory retrieval, as a general process and collapsed across all memory retrieval trials (15 memory trials per run), compared to the control counting task (generalised episodic memory retrieval); and, (B) whether focal hippocampal subfield damage impaired the ability to represent specific episodic memories (specific episodic memory retrieval). Each memory cell contains five trials per run of that specific memory (denoted as Trial (T) 1, T2, T3, T4, and T5). Each of the 30 trials (15 memory and 15 counting) per run were then subjected to trial-by-trial correlation analyses. Counting trials are not shown in (B) for simplicity. Model matrices were constructed to feature a value of 1 in hypothesized high-dissimilarity cells and a value of 0 in hypothesized low-dissimilarity cells.

For the first model (Figure 5A), significant model fit was observed throughout the AMN in both groups (see Supplementary data), except for a significant between-group difference in the left angular cortex (amnesic group mean Kendall τ: 0.395±0.078, control group mean Kendall τ: 0.456±0.47; *t*_(34)_ = 2.830; *p* = 0.008, Cohen’s d = 0.943).

**Figure 5.**
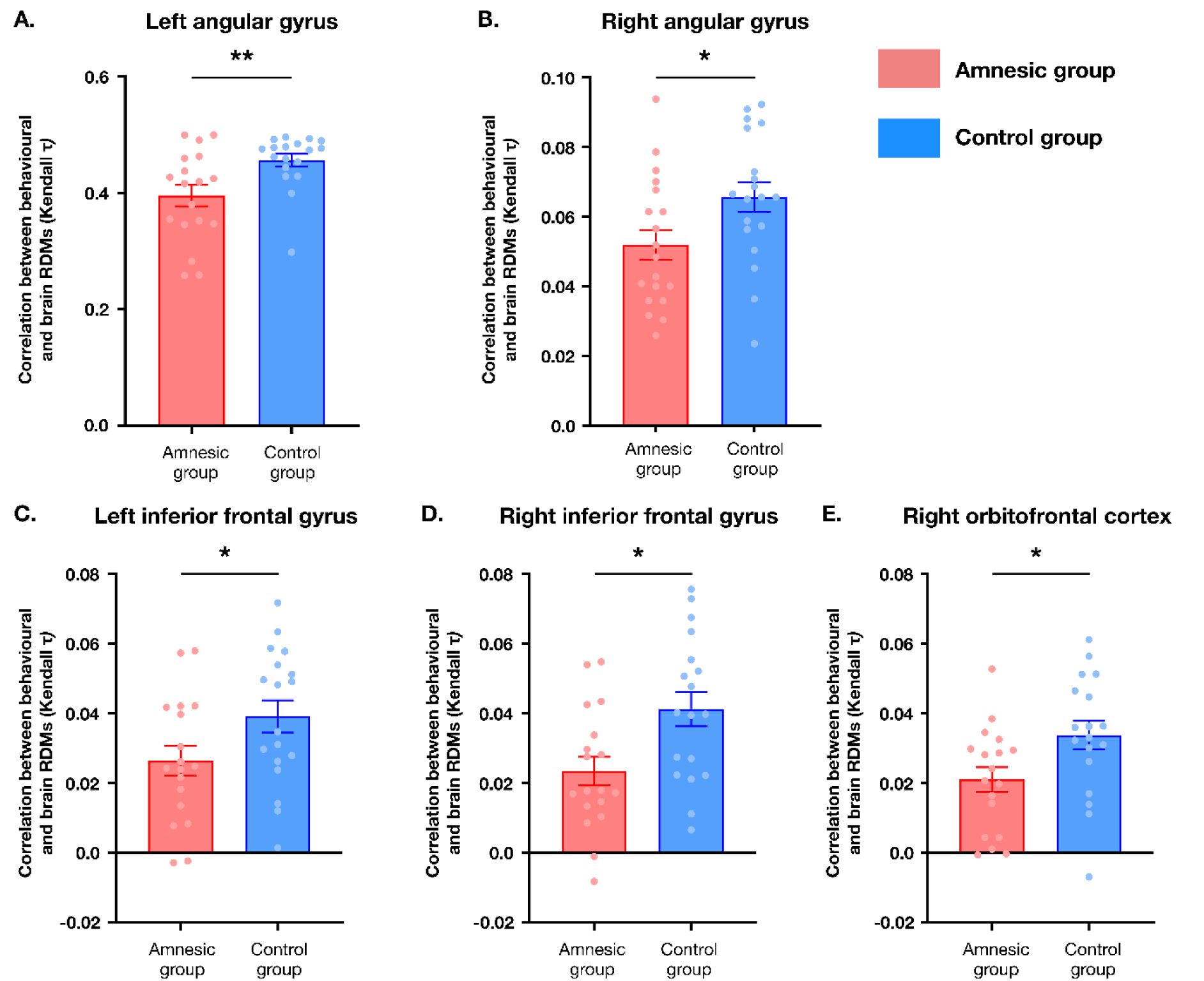
Hippocampal subfield damage selectively impaired the autobiographical memory network. RDMs were used to conduct two model fit analyses: (1) a model reflecting general episodic memory retrieval, independently of the specific episodic memory being remembered; and, (2) a model focused on the retrieval of specific episodic memories (see Figure 4). As detailed in Supplementary data, hippocampal subfield pathology did not prevent significant model fits for both hypotheses matrices across the majority of tested cortical regions. Group differences were found, however, in (A) the left angular gyrus for the model specifying representational content during general episodic memory retrieval (amnesic group mean Kendall τ: 0.395±0.078, control group mean Kendall τ: 0.456±0.47; *t*_(34)_ = 2.830; *p* = 0.008, Cohen’s d = 0.943); and in (B) the right angular cortex (amnesic group mean Kendall τ: 0.0519±0.0045; control group mean Kendall τ: 0.0656±0.0045; *t*_(34)_ = 2.141; *p* = 0.040, Cohen’s d = 0.714), (C) left inferior frontal gyrus (amnesic group mean Kendall τ: 0.0364±0.0046; control group mean Kendall τ: 0.0391±0.0046; *t*_(34)_ = 2.038; *p* = 0.049, Cohen’s d = 0.679), (D) right inferior frontal gyrus (amnesic group mean Kendall τ: 0.0233±0.0041; control group mean Kendall τ: 0.0412±0.0049; *t*_(34)_ = 2.778; *p* = 0.009, Cohen’s d = 0.926), and (E) right orbitofrontal cortex (amnesic group mean Kendall τ: 0.00210±0.0035; control group mean Kendall τ: 0.0337±0.0041; *t*_(34)_ = 2.357; *p* = 0.024, Cohen’s d = 0.786) for the hypothesis model for specific episodic memories. Error bars correspond to the standard error of the mean (n = 18). * denotes *p*<0.05; ** denotes *p*<0.01.

For the second model (mnemonic discriminability; Figure 5B-E), significant model fit was observed throughout the AMN network in both groups (see Supplementary data), except for significant between-group differences in the right angular cortex (rANG; amnesic group mean Kendall τ: 0.0519±0.0045; control group mean Kendall τ: 0.0656±0.0045; *t*_(34)_ = 2.141; *p* = 0.040, Cohen’s d = 0.714), left inferior frontal gyrus (lIFG; amnesic group mean Kendall τ: 0.0364±0.0046; control group mean Kendall τ: 0.0391±0.0046; *t*_(34)_ = 2.038; *p* = 0.049, Cohen’s d = 0.679), right inferior frontal gyrus (rIFG; amnesic group mean Kendall τ: 0.0233±0.0041; control group mean Kendall τ: 0.0412±0.0049; *t*_(34)_ = 2.778; p = 0.009, Cohen’s d = 0.926), and the right orbitofrontal cortex (rOFC; amnesic group mean Kendall τ: 0.00210±0.0035; control group mean Kendall τ: 0.0337±0.0041; *t*_(34)_ = 2.357; p = 0.024, Cohen’s d = 0.786).

The fMRI data were also assessed for between-group differences in the mnemonic representational stability for the specific memories arising from the first-level Fisher-transformed Pearson dissimilarity scores (as a proxy of mnemonic representational stability; Figure 6). An omnibus 2 (group: amnesic group, controls) x 3 (memory: three experimental episodic memories) ANOVA was conducted. Mauchly’s test demonstrated that the assumption of sphericity had not been violated. There were significant main effects of group for the rANG (amnesic group mean: 0.76±0.03; control group mean: 0.68±0.02; *F*_(1,34)_ = 6.156, *p* = 0.018, η^2^ = 0.153), the rIFG (amnesic group mean: 0.88±0.02; control group mean: 0.81±0.02; *F*_(1,34)_ = 9.129, *p* = 0.005, η^2^ = 0.213), and the rOFC (amnesic group mean: 0.89±0.02; control group mean: 0.83±0.02; *F*_(1,34)_ = 6.969, *p* = 0.012, η^2^ = 0.170).

**Figure 6.**
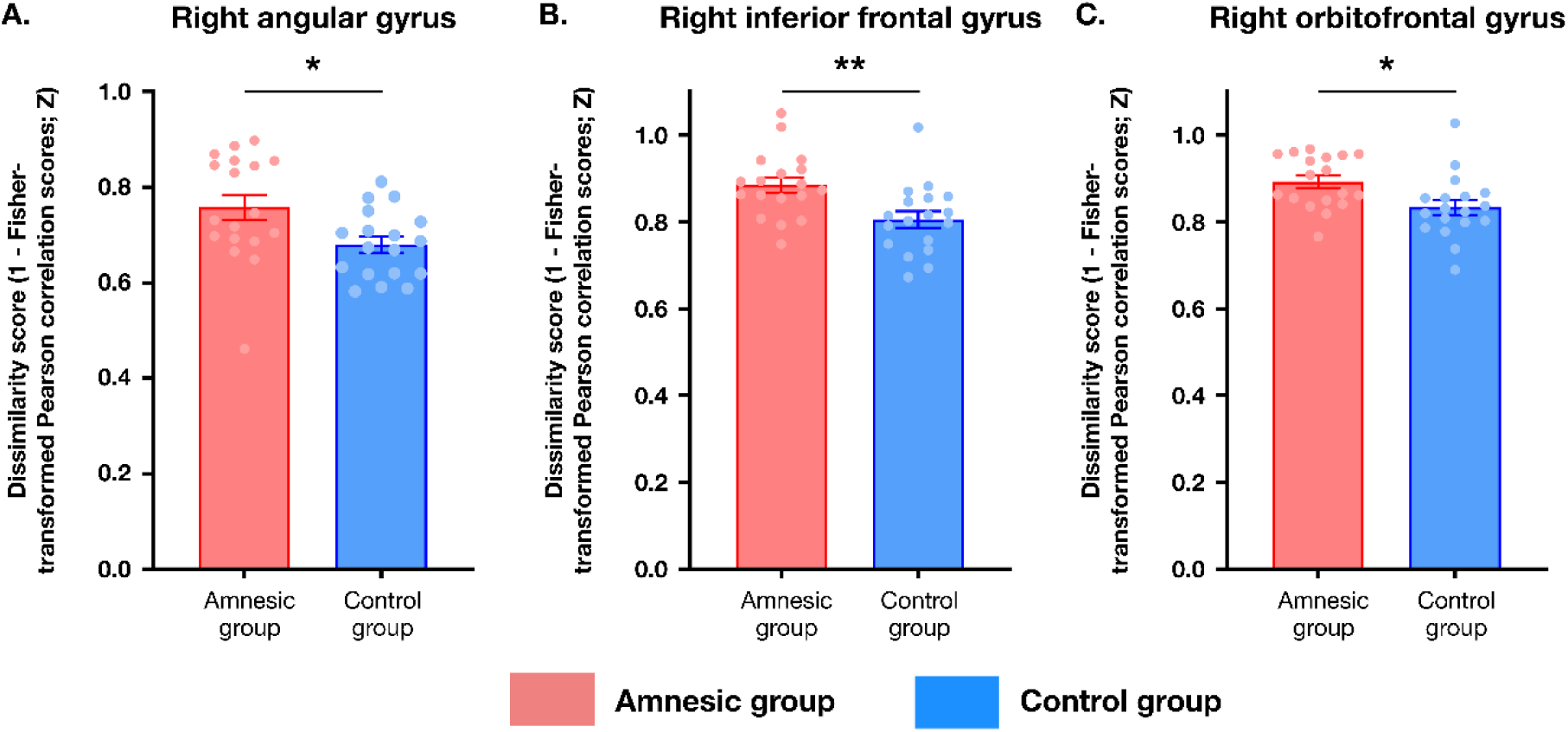
Hippocampal subfield damage was associated with decreased mnemonic stability for specific episodic memories. To investigate these between-group level differences in specific memory representation further, we tested for between-group differences in the Fisher-transformed Pearson correlation scores (as a measure of mnemonic representational stability; lower dissimilarity scores indicate greater voxel population stability for each retrieval trial) corresponding to specific memories from the first-level analysis of the RDMs. This demonstrated that there was a significant main effect of group (n = 18; but no main effect of memory or an interaction between the group and memory) in these Fisher-transformed Pearson correlation scores for the (A) right angular gyrus (amnesic group mean: 0.76±0.03; control group mean: 0.68±0.02; *F*_(1,34)_ = 6.156, *p* = 0.018, η^2^ = 0.153); (B) right inferior frontal gyrus (amnesic group mean: 0.88±0.02; control group mean: 0.81±0.02; *F*_(1,34)_ = 9.129, *p*=0.005, η^2^ = 0.213); and, (C) right orbitofrontal cortex (amnesic group mean: 0.89±0.02; control group mean: 0.83±0.02; *F*_(1,34)_ = 6.969, *p* = 0.012, η^2^ = 0.170). No main effect of memory was seen for the left inferior frontal gyrus (amnesic group mean: 0.87±0.02; control group mean: 0.83±0.02; *F*_(1,34)_ = 2.753, *p* = 0.106, η^2^ = 0.075). * denotes *p*<0.05; ** denotes *p*<0.01.

Importantly, no main effect of memory [rANG: *F*_(2,68)_ = 1.257, *p* = 0.291, η^2^ = 0.036; rIFG: *F*_(2,68)_ = 0.889, *p* = 0.416, η^2^ = 0.025; rOFC: *F*_(2,68)_ = 2.411, *p* = 0.097, η^2^ = 0.066] nor significant group by memory interactions [rANG: *F*_(2,68)_ = 1.720, *p* = 0.187, η^2^ = 0.048; rIFG: *F*_(2,68)_ = 0.562, *p* = 0.573, η^2^ = 0.016; rOFC: *F*_(2,68)_ = 1.141, *p* = 0.326, η^2^ = 0.032] were observed. For the lIFG, the ANOVA revealed no significant main effects of group (amnesic group mean: 0.87±0.02; control group mean: 0.83±0.02; *F*_(1,34)_ = 2.753, *p* = 0.106, η^2^ = 0.075) and memory (*F*_(2,68)_ = 0.056, *p* = 0.945, η^2^ = 0.002), and the group by memory interaction was also non-significant (*F*_(2,68)_ = 0.18, *p* = 0.982, η^2^ = 0.001).

### CA23 but not CA1 volume predicted the representational stability of episodic memories

Next, we sought to investigate whether there was a causal link between the variability in total CA23 and/or total CA1 volumes and the two hypothesis model fits (as measured by Kendall τ for general and specific episodic memory retrieval), as well as the stability in the mnemonic representational similarity across trials (in this case, the averaged Fisher-transformed Pearson dissimilarity scores arising from the first-level analysis). Stepwise regressions were undertaken with total CA23 and total CA1 volumes as the independent variables and Kendall τ and averaged Fisher-transformed Pearson correlation values as the dependent variables.

For general episodic memory retrieval, a significant model was identified through stepwise regression (*F*_(1,34)_ = 6.452), where total CA23 volume emerged as a selective predictor of episodic memory retrieval model fit in the lANG (*t* = 2.540, *p* = 0.016, R² = 0.159, β₁ = 0.399; Figure 7A). Total CA1 volume was excluded from this model even when the order of entry was reversed.

**Figure 7.**
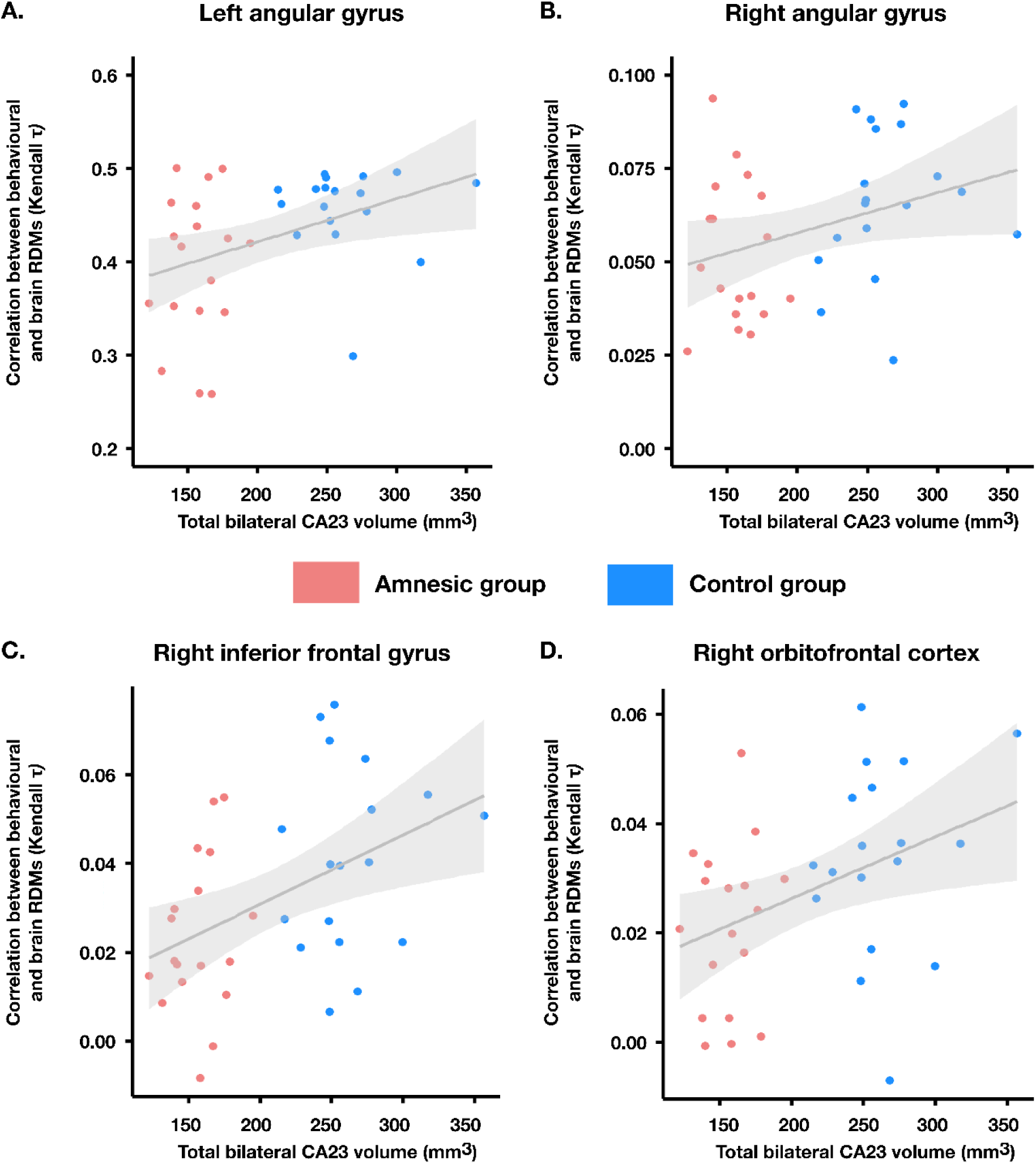
CA23 volume predicted RDM model fits across both groups. On the basis of the causal link between the CA23 volume and cumulative episodic memory performance, we sought to investigate whether there was also a causal link between the variability in total CA23 and/or total CA1 volumes and the two hypothesis model fits (as measured by Kendall τ; total population n = 36). Stepwise regressions were undertaken with total CA23 and total CA1 volumes as the independent variables and Kendall τ correlation values as the dependent variables. (A) For the hypothesis matrix specifying general episodic memory retrieval in the left angular gyrus, a significant model was identified through stepwise regression (*F*_(1,34)_ = 6.452), where total CA23 volume emerged as a selective predictor of episodic memory recall (*t* = 2.540, *p* = 0.016, R² = 0.159, β₁ = 0.399). Total CA1 volume was excluded from this model, even when the order of entry was reversed (*t* = 0.078, *p* = 0.938). When considering the hypothesis matrix specifying individual memories, significant models were found in for the (B) right angular gyrus (*F*_(1,34)_ = 5.076, *p* = 0.031); (C) right inferior frontal gyrus (*F*_(1,34)_ = 8.605, *p* = 0.006); and, (D) right orbitofrontal cortex (*F*_(1,34)_ = 6.354, *p* = 0.017). All of these models were then associated with total CA23 volume selectively predicting the hypothesis model fit (rANG: *t* = 2.253, *p* = 0.031, R^2^ = 0.130, β₁ = 0.360; rIFG: *t* = 2.930, *p* = 0.006, R^2^ = 0.202, β₁ = -0.449; rOFC: *t* = 2.521, *p* = 0.017, R^2^ = 0.157, β₁ = 0.397). In all cases, total CA1 volume was excluded from the model even when the order of entry into the model was reversed (rANG: *t* = 0.671, *p* = 0.418; rIFG: *t* = - 0.606, *p* = 0.549; rOFC: *t* = -0.403, *p* = 0.689). Graphs show scatter plots (with best-fitting linear regression line) illustrating the relationship between the magnitude of total CA23 volume as an independent variable selectively predicting the model fit scores (total population n = 18).

For specific memory retrieval, significant models (Figure 7B-D) were found in for the rANG (*F*_(1,34)_ = 5.076, *p* = 0.031), rIFG (*F*_(1,34)_ = 8.605, *p* = 0.006), and rOFC (*F*_(1,34)_ = 6.354, *p* = 0.017). Crucially, total CA23 volume selectively predicted the hypothesis model fit (rANG: *t* = 2.253, *p* = 0.031, R^2^ = 0.130, β₁ = 0.360; rIFG: *t* = 2.930, *p* = 0.006, R^2^ = 0.202, β₁ = -0.449; rOFC: *t* = 2.521, *p* = 0.017, R^2^ = 0.157, β₁ = 0.397). In all cases, total CA1 volume was excluded from the model, even when the order of entry into the model was reversed (rANG: *t* = 0.671, *p* = 0.418; rIFG: *t* = -0.606, *p* = 0.549; rOFC: *t* = -0.403, *p* = 0.689). A non-significant stepwise regression model was found for lIFG (*F*_(2,35)_ = 2.596, *p* = 0.090).

When considering the representational stability (where lower dissimilarity scores signify greater trial-by-trial similarity in the neuronal populations supporting those memories, Figure 8A-C), significant models were found in all three regions rANG: *F*_(1,34)_ = 6.076, *p* = 0.019; rIFG: *F*_(1,34)_ = 10.877, *p* = 0.002; and, rOFC: *F*_(1,34)_ = 6.354, *p* = 0.017). Notably, the variability in CA23 volume predicted the representational in all three regions (rANG: *t* = -2.465, *p* = 0.019, R^2^ = 0.152, β₁= - 0.389; rIFG: *t* = -3.298, *p* = 0.002, R^2^ = 0.242, β₁ = -0.492; rOFC: *t* = -3.382, *p* = 0.002, R^2^ = 0.252, β₁ = -0.502). Once again, total CA1 volume was excluded from the model, even when the order of entry was reversed (rANG: *t* = 1.193, *p* = 0.241; rIFG: *t* = 1.626, *p* = 0.113; rOFC: *t* = 0.449, *p* = 0.656).

**Figure 8.**
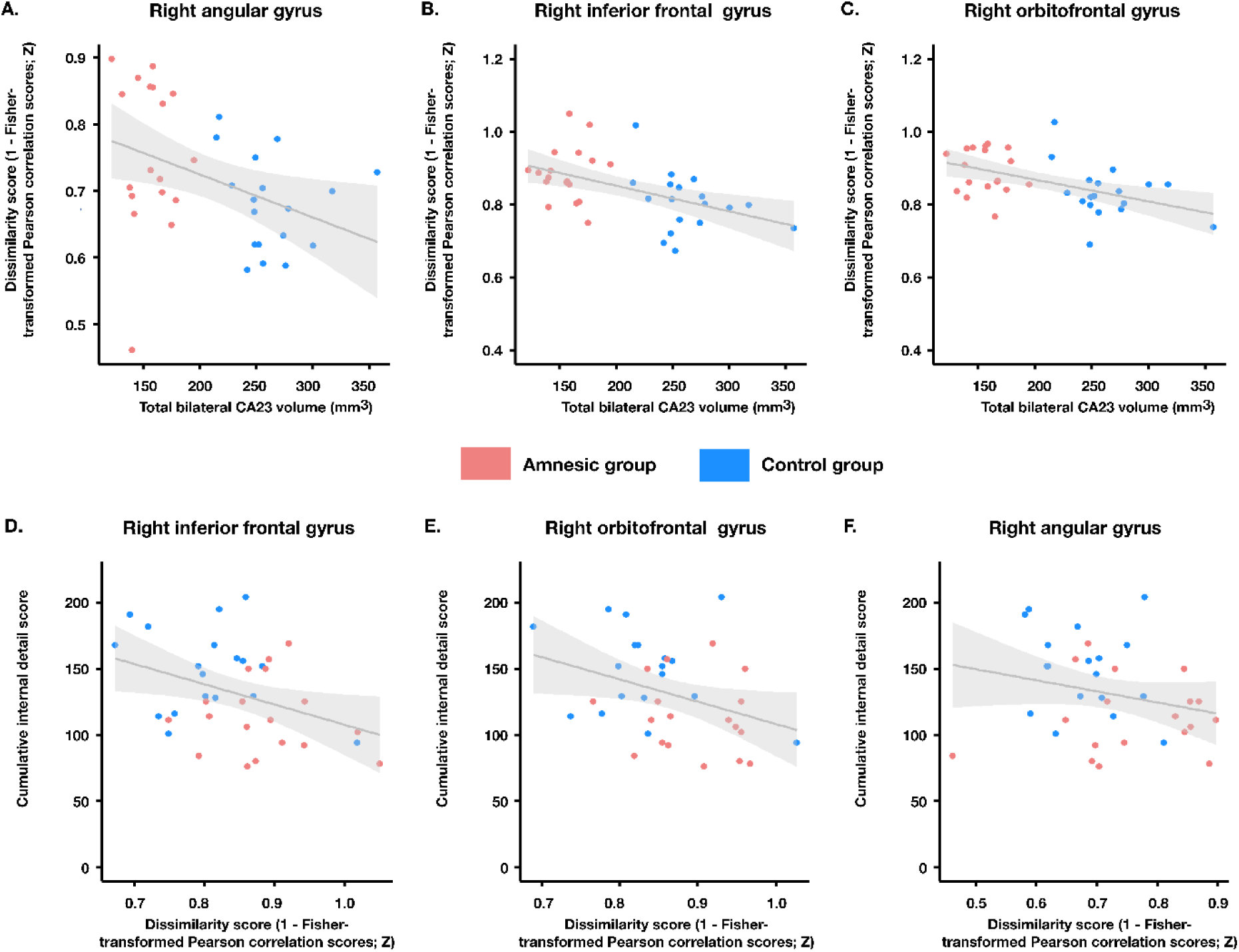
CA23 volume predicted episodic memory representational stability and representational stability predicted the episodic details associated with personal events. When considering relationship between the total CA23 volumes and the averaged dissimilarity scores (where lower scores signify lower trial-by-trial dissimilarity in the voxel populations supporting those memories; calculated by 1 - Fisher-transformed Pearson correlation scores), significant models were once again discovered in all three regions: (A) right angular gyrus (F_(1,34)_=6.076, p=0.019); (B) right inferior frontal gyrus (*F*_(1,34)_ = 10.877, *p* = 0.002); and, (C) right orbitofrontal cortex (*F*_(1,34)_ = 6.354, *p* = 0.017). The variability in CA23 volume also predicted the averaged Fisher-transformed Pearson scores in all three regions (rANG: *t* = -2.465, *p* = 0.019, R^2^ = 0.152, β₁ = -0.389; rIFG: *t* = -3.298, *p* = 0.002, R^2^ = 0.242, β₁ = -0.492; rOFC: *t* = -3.382, *p* = 0.002, R^2^=0.252, β₁ = -0.502). Once again, total CA1 volume was excluded from the model even when the order of entry was reversed (rANG: *t* = 1.193, *p* = 0.241; rIFG: *t* = 1.626, *p* = 0.113; rOFC: *t* = 0.449, *p* = 0.656). Graphs show scatter plots (with best-fitting linear regression line) illustrating the relationship between the magnitude of total CA23 volume as an independent variable selectively predicting the averaged Fisher-transformed Pearson scores for the three experimental memories. Averaged mnemonic representational dissimilarity scores across recall trials were used as the independent variable in a linear regression with cumulative internal (episodic) detail score as the dependent variable. We found that the similarity of episodic memory representations selectively predicted internal (episodic) memory score in the (D) rIFG (*t* = 2.230, *p* = 0.032, R^2^ = 0.128, β₁ = 0.357) and (E) rOFC (*t* = -2.189, *p* = 0.036, R^2^ = 0.124, β₁ = -0.352), but not (F) rANG (*t* = -1.452, *p* = 0.156, R^2^ = 0.058, β₁ = -0.242). Graphs show scatter plots (with best-fitting linear regression line) illustrating the relationship between the magnitude of the dissimilarity scores for the three episodic memories (per participant) as the independent variable selectively predicting the cumulative internal (episodic) scores for the three experimental memories (total population n = 18).

### Representational stability predicted the remembering of episodic (internal) details associated with personal events

Finally, we wanted to test whether there was a causal relationship between the stability of episodic memory representations (using the first-level Pearson correlation dissimilarity scores) and internal (episodic) memory performance for the experimental memories. The results revealed that the stability of episodic memory representations selectively predicted internal (episodic) memory score in performance in the rIFG (*t* = 2.230, *p* = 0.032, R^2^ = 0.128, β₁ = 0.357) and rOFC (*t* = -2.189, *p* = 0.036, R^2^ = 0.124, β₁ = -0.352), but not rANG (*t* = -1.452, *p* = 0.156, R^2^ = 0.058, β₁ = -0.242; see Figure 8D-F).

### Summary

The results supported and elaborated our *a priori* hypotheses for the fMRI study. Specifically, the group with amnesia showed: (1) significantly reduced mnemonic distinctiveness for (2) regions within the AMN with close connections to the hippocampus. In addition, (3) the stability of episodic memory representations in the rIFG and rOFC, but not rANG selectively predicted internal (episodic) memory score and (4) total CA23 volume predicted general episodic memory retrieval in the lANG, and the retrieval of specific episodic memories and their stability in the rANG, rIFG, and rOFC.

## DISCUSSION

A large-scale hippocampal-neocortical network supports episodic memory for autobiographical events and experiences,^3,31,88–92^ yet it is unknown how human hippocampal damage affects the representational content in the nodes within this hippocampal-neocortical network. To address this question, we investigated the effects of human hippocampal damage on the representational content of pre-morbid, recently acquired autobiographical memories during retrieval. Results from structural brain neuroimaging and cognitive phenotyping using the AI demonstrated that the hippocampal damage was bilateral and focal and associated with reduced episodic detail retrieval, with variability in total CA23 volume predicting the amount of remembered episodic detail. These results replicate previous findings in an entirely new cohort of individuals with amnesia.^31,37^ In addition, we found that focal hippocampal pathology was associated with a mild impairment on standardised assessments of anterograde memory, while the group differences in behaviour reflected specific memory deficits rather than generalised cognitive dysfunction.^31,37^

Results from fMRI RDM-based analyses revealed that, in line with our first and second hypotheses, hippocampal damage was associated with between-group differences in brain-derived RDM model fit scores, revealing reduced mnemonic dissimilarity (or distinctiveness) in the group with hippocampal amnesia across a limited number of cortical regions within the AMN. Specifically, between-group differences in the representation of the episodic memory retrieval state (versus the baseline counting state) were limited to the lANG, whereas differences in the representation of specific episodic memories were found in the rANG, rOFC, lIFG and rIFG. This circumscribed effect is notable, given that the hippocampus is an important hub region,^78–81^ and suggests that hippocampal damage can have a limited, rather than widespread, impact on memory representations in cortical regions. These more limited effects on cortical network nodes align with previous graph-theoretic based analyses of multiple resting-state networks, which showed that the disruptive effects of focal CA3 damage led to limited effects on network topology.^31^ Importantly, hippocampal subfield volume predicted the magnitude of dissimilarity between specific episodic memories, with the variability in model fit across all regions specifically predicted only by total CA23 volumes. Hence, we show that CA23 not only has a critical role in supporting the retrieval of episodic memories, but also in supporting the maximal dissimilarity in the neocortical neural representation of competing memories.

Having discovered the impact of hippocampal damage on the representation of dissimilar episodic memories, we tested our third hypothesis by examining the effects of hippocampal pathology on the stability of specific mnemonic representation,^69^ finding that, compared with the control group, the group with hippocampal amnesia had increased trial-by-trial dissimilarity in the voxel patterns within the rANG, rOFC, and rIFG; in addition, the variability in these values was predicted by total CA23 volume. Crucially, we discovered that lower trial-by-trial dissimilarity in the rOFC and rIFG was predictive of the retrieval of greater episodic memory detail. In support of our fourth hypothesis, reduced CA23 volumes were therefore not only associated with less differentiated neural representations of specific episodic memories, but also an impairment in the stability of the trial-by-trial voxel populations that supported those episodic memories.

Both distinctiveness and stability are crucial for memory performance, and have been described as separate properties of neural representations.^69^ These effects align with the dual role of CA3 supporting computations that make overlapping experiences more distinct (pattern separation).^77^ and those that code relationships across items to increase overlap (pattern completion).^38,82,89,93–96^ In healthy adults, CA3 volume has previously been found to scale linearly with the degree of differentiation between neuronal voxel populations within the hippocampus for specific episodic memories, which contributes to efficient pattern separation.^64^ Reduced CA23 volumes might therefore be associated with less efficient pattern separation,^76^ leading to reduced dissimilarity between competing episodic memories within the hippocampus, whereas voxel population instability could indicate compromised pattern completion. Conceivably, decreased mnemonic overlap within CA3 is associated with increased mnemonic competition within the neocortex, potentially leading to reduced episodic memory re-experiencing.^64^ Here, we demonstrated that the stability of those neocortical voxel ensembles was linked with CA23 volume loss, and this instability was indirectly associated with the retrieval of less re-experiential and detailed episodic memories. CA23 pathology may, therefore, disrupt the balance between pattern separation and completion computations that underscores successful episodic memory retrieval.

Next, we consider in greater detail how reduced dissimilarity in the affected nodes of the AMN (lANG, rANG, lIFG, rIFG, and rOFC) is likely to be translated into impaired episodic memory. In the angular gyrus (ANG), we found that lANG was implicated in the state of episodic memory retrieval, whereas the rANG was implicated in the representation of specific episodic memories. The angular gyrus maintains extensive connections with multiple neocortical regions and plays a crucial role in memory tasks that require detailed recollection or complex integration of information across sensory modalities.^5,97–101^ Emerging evidence suggests distinct functional roles for the left and right hemispheric components. The lANG is functionally connected with the dorsomedial PFC subsystem^102^ of the default mode network (DN),^103^ a subsystem that has been implicated in subjective elements of confident recollection rather than the retrieval of the objective context.^102,104^ The lANG may, in alignment with our RDM-based results reported here, support the generalised and subjective aspects of recollective confidence and re-experiential richness,^105^ rather than the objective recollection of details forming that specific event.^102,106^ In contrast, the rANG appears to have greater functional connectivity with components of the MTL subsystem of the DN during episodic memory retrieval.^104^ The MTL subsystem is implicated in the retrieval of source memories^104^ and the retrieval of specific temporal order,^107^ with the rANG having a specific role in supporting the objective retrieval of specific episodic memory details.^102^ Our results would seem to corroborate a putative role of the rANG in representing specific episodic mnemonic details, given that hippocampal damage in the amnesic group was associated with a diminished ability to represent distinct episodic memories. Hence, it would appear that diminished mnemonic distinctiveness impaired both the subjective experience of episodic memories supported by the lANG and the remembering of specific episodic details mediated by the rANG. Studying specific subdivisions within the angular gyrus may reveal new insights into the mechanisms behind episodic memory impairments.

Hippocampal pathology also impaired specific mnemonic representation in the lIFG, rIFG, and rOFC. These findings align with previous studies that have shown the IFG is crucial for episodic memory,^108–112^ and has also been implicated in other cognitive control processes including task monitoring,^113^ cognitive control,^114–116^ representing imagined stimuli,^115,117,118^ suppressing irrelevant memories,^119^ and sensitivity to cue specification.^120^

The rOFC was also implicated in the representation of specific episodic memories, which is in line with previous studies of episodic learning^121–124^ and retrieval,^125^ where univariate BOLD signal increase within the OFC have been linked to subsequent memory performance.^121^ The hippocampus and OFC interact closely during episodic memory tasks, with the hippocampus providing crucial contextual information to the OFC to facilitate the integration of episodic details within broader cognitive frameworks.^126,127^ Therefore, hippocampal damage may disrupt the integration of context-specific memory representations to the OFC,^128^ thereby impeding the formation of a coherent cognitive representation of the memory.^128–130^ Support for this contention comes from OFC lesion studies that have revealed difficulty in distinguishing between relevant and irrelevant memory traces, alongside content misattribution.^124,131–133^ The OFC is recruited after other components of the AMN, which suggests that it exerts important evaluative and integrative functions central to memory-based decision-making processes,^125^ thereby functioning to mediate between specific memories.^134^ In summary, reduced episodic mnemonic discrimination within IFG and OFC was associated with less fine-grained episodic detail, perhaps due an inability to represent distinct and/or stable memories that are required for supporting these later-stage monitoring or evaluative processes.

More broadly, the angular gyrus, OFC, and IFG form part of the frontoparietal control network (FPCN),^103,135,136^ which contributes to executive control and guiding goal-based actions.^137^ The FPCN is closely connected with the DN and the dorsal attention network (DAN).^138^ The FPCN can be fractionated into two distinct subsystems, using resting state-fMRI: (1) FPCN subsystem A (FPCN_A_; comprised in part by the OFC and angular gyri), which is closely associated with the DN; and, (2) FPCN subsystem B (FPCN_B_; comprised, in part, by the IFG), which is closely associated with the DAN. Functionally, the FPCN_A_ regions are associated with tasks requiring introspective attention away from perceptual input^139,140^ and episodic memory.^141–143^ Given that the DN facilitates the integration of conceptual-associative knowledge into current thought and perception^144–146^, hippocampal damage could potentially destabilise the representation of internally generated mnemonic re-experiencing. FPCN_B_ regions appear closely related to cognitive control by encoding task-relevant information including tasks rules and their relationship to expected reward outcomes,^135,147–150^ as well as through top-down modulation of the DAN.^151,152^ Hippocampal damage may impair the ability for the FPCN_B_ to utilise distinct and/or stable mnemonic information in order to maintain top-down attention on specific episodic memory retrieval.

Importantly, our data show that hippocampal damage did not eliminate specific mnemonic representations across other nodes within the AMN. This is not unexpected, especially given that selective hippocampal damage impairs certain aspects of memory retrieval, yet it does not prevent remembering all details related to specific episodic memories.^22,23,25–28,30,31,37,77,153^ Some accounts posit that the hippocampus is required for episodic retrieval where the memory has spatial detail and context-specific content.^3,5,40,41,154^ Accordingly, hippocampal damage could render episodic memories ‘gist-like’ in nature yet remain discrete events.^3,5,40,41,154^ Given that the retrieval of richly re-experiential memories requires coordination of multiple cortical regions,^3,88–92^ the retrieval of the gist or context of a memory could be hippocampal-independent^3,5,40,41,154^. Under such an account, the hippocampus is critical for integrating sensory, perceptual, and conceptual information into a cohesive experience.^1–5^. Incorporating this account into what is known about CA3 mediated computations may explain our findings in terms of residual functionality. Specifically, the hippocampal pathology may prevent the representation of distinct and/or stable memories, resulting in the observed episodic impairments, but still permit some degree of mnemonic discriminability to support memory retrieval. This interpretation aligns with the idea that hippocampal damage can affect the richness and detail of memories rather than completely abolishing episodic remembering, and it provides a potential mechanism for the observed preservation of gist-like memory retrieval in the context of hippocampal pathology.

We also replicated previous findings on the anatomical effects of the LGI1-LE aetiology, thereby confirming that the damage was limited to bilateral hippocampal regions. A prior study involving eighteen individuals with the same, single aetiology reported bilateral focal CA3 volume loss (∼28% volume loss compared to controls)^31,37^. In the current study, there was an almost 40% reduction in CA23 volume compared to the control group. One source of this notional difference is that CA2 and CA3 were not separately delineated due to technical reasons, including the use of lower field-strength MRI (3.0-Tesla vs 7.0-Tesla) and differences in other acquisition parameters, which led to differences in biophysical appearance of the hippocampal subfields. Delineation of the hippocampal subfields in the current study was also performed with a semi-automated segmentation pipeline on the 3.0-Tesla dataset, with manual validation versus a protocol limited to manual segmentation on the 7.0-Tesla dataset. Contrary to previous results, the current cohort also demonstrated significant CA1 atrophy as well. The current data, however, remain comparable with those previous results. The previous non-significant CA1 result was observed in the context of a Holm-Bonferroni corrected alpha-criterion,^31,37^ yet the proportion of volume loss in CA1 reported here from 18 individuals with the same presentation and aetiology is similar (18% in the current study and 15% previously). CA1 atrophy is not necessarily unexpected as there is evidence of neuronal loss in both CA3 and CA1 following seizures in homozygous LGI1 knockout mice.^155^ Further data will be needed to adjudicate as to whether significant volume loss in CA1 is a feature of LGI1-LE. Notably, our data also demonstrated that the variability in total CA1 volumes did not have any predictive power across the behavioural and imaging data compared to total CA23 volumes. In line with prior studies, we also found no supra-threshold clusters of grey matter volume loss were detected in the hippocampal amnesia compared to the controls. Together, the hippocampal segmentation and whole-brain data align with those reported by other laboratories.^31,37,48,54,55,156^

In terms of theoretical significance for accounts of episodic memory, our results accord with recent reconstructive models of hippocampal function,^1–5,41^ whereby hippocampal pathology did not prevent the representation of discrete autobiographical events, instead it impaired the cortical dissimilarity and stability of specific autobiographical memories in only a small subset of AMN nodes. We were limited by voxel size in this study, but it would be critical to assess mnemonic dissimilarity within individual hippocampal subfields to understand whether a failure of pattern separation occurs within CA23. Moreover, to ensure hippocampal involvement, we specifically chose to evaluate memories from a recent time point that, according to multiple models of hippocampal function, should still rely on hippocampal processing for retrieval.^4,5,12,41,44,157–159^ Accordingly, the current data support theories that place the hippocampus as a central coordinating hub within the AMN, with hippocampal damage altering mnemonic representations in distal cortical regions.^1–5,21^ In future work, it will be crucial to establish whether memories arising from more distant timepoints have the same pattern of decreased mnemonic dissimilarity as these hippocampal-dependent memories^12^.

To conclude, our results have addressed a key question: How does focal hippocampal damage, associated with episodic amnesia, affect the representational content and brain network that underlies autobiographical memories? We found that hippocampal pathology prevented the distinct and stable representation of episodic memories in a sparse number of cortical components of the AMN, albeit ones with critical functions to episodic memory retrieval. We were able to relate the stability of voxel activity patterns across each memory retrieval trial to the amount of episodic memory details retrieved, thereby providing, for the first time, the first direct neural correlate between focal hippocampal dysfunction, altered mnemonic representational content, and episodic amnesia. Future research should explore the temporal dynamics of cortical representations during memory formation and retrieval to better understand how hippocampal-cortical interactions enable flexible event component reuse. In addition, investigating how different cortico-hippocampal networks represent specific event components during online experience and memory retrieval could enhance our understanding of how the brain scaffolds memory for various high-level event elements.

## Methods

### Participants

All participants in the group with hippocampal amnesia had chronic amnesia induced by a single-aetiology: leucine-rich glycine-inactivated-1 limbic encephalitis (LGI1-LE).^160^ All participants were seizure-free at the time of testing and considered clinically stable by their treating neurologist (TM, MZ, AH, TP, EC). The group with hippocampal amnesia was comprised of 18 participants (mean and S.E.M. age: 58.1±2.33 years, male = 13). Eighteen healthy age- and education-matched controls were recruited (58.1± 2.33 years, male = 13) to the control group. These control participants had no history of cognitive, psychiatric, or neurological illness. All participants had normal or corrected-to-normal vision. All participants were fluent, native English speakers. No significant differences were found between the groups in terms of age or years-of-education. LGI1-LE is a typically monophasic illness with no ongoing change in postmorbid cognition expected;^31,37,87^. Accordingly, none of the participants with amnesia had clinical evidence of a relapse at any time surrounding the experimental period. All participants with amnesia were otherwise self-reported as healthy, with no evidence of secondary gain or active psychopathology. Informed written consent was obtained from all participants for all procedures and for consent to publish, in accordance with the terms of approval granted by a national research ethics committee (22/NW/0088) and the principles expressed in the Declaration of Helsinki. Care was taken to ensure that all aspects of this experiment (including task design, administration, and task comprehension) were undertaken to provide maximal retrieval support and facilitate task comprehension in both groups.

### Neuropsychological assessment

Neuropsychological assessment was conducted in both groups using the following standardized neuropsychological subtests: *Verbal and visual intelligence:* Wechsler Abbreviated Scale of Intelligence (WASI) – Similarities and Matrix Reasoning;^161^ *Premorbid intelligence:* Test of Premorbid Functioning – UK Version;^162^ *Verbal memory:* Logical memory I and II from Wechsler Memory Scale–III (WMS-III);^163^ *Visual memory:* Rey complex figure – Immediate-Recall and Delayed-Recall;^164^ *Recognition memory*: Recognition Memory Test – Words and Faces;^165^ *Sustained Attention*: Test of Everyday Attention – Map Search;^166^ *Language*: Graded Naming Test;^167^ Letter Fluency and Category Fluency from the Verbal Fluency from Delis-Kaplan Executive Function System (D-KEFS);^168^ *Executive function:* Category Switching from the Verbal Fluency Test, Number-Letter Switching from the Trail Making Test from Delis-Kaplan Executive Function System (D-KEFS);^168^ and, WMS-III – Digit Span;^163^ *Attentional Switching / Cognitive Flexibility:* Visual Elevator from Test of Everyday Attention;^166^ and, Number-Letter Switching from the Trail Making Test from the Delis-Kaplan Executive Function System (D-KEFS);^168^ *Visuomotor skills:* Visual Scanning, Number Sequencing, Letter Sequencing, and Motor Speed from the Trail Making Test from Delis-Kaplan Executive Function System (D-KEFS;)^168^ and *Visuoconstruction skills*: Rey complex figure – Copy.^164^ Scores on the standardized neuropsychological tests were first transformed into age-corrected standard values, where available, and then transformed into *Z*-scores. One participant in the amnesic group was unable to complete the premorbid IQ (TOPF) test due to severe dyslexia.

### Neuropsychology statistical analyses

The data from the neuropsychological assessments were analysed in two ways: (1) two-sample *t*-tests (two-tailed) conducted to determine whether amnesic group performance was significantly different from normative data (with a mean of 0 and standard deviation of 0.3); and, (2) a between-groups independent-sample *t*-test (two-tailed) to determine where group level performance was significantly different between the two experimental groups.

### Autobiographical interview

As LGI1-LE is a discrete, autoimmune condition, symptom-onset can be reliably identified; therefore, we were able to acquire specific autobiographical memories that occurred two-to-five years before the onset of the illness, a period where autobiographical memories are hypothesised to become consolidated across the autobiographical memory network.^12,44,157–159^ The monophasic nature of autoimmune encephalitis suggests that hippocampal function was intact in these participants during the encoding of memories before the disease onset. Impaired retrograde memory, therefore, probably reflects the disruption of hippocampal-mediated retrieval mechanisms rather than anterograde difficulties with encoding and consolidation. Memories were obtained from the 18 participants in the hippocampal amnesia and 18 participants in the control group 14 days prior to the scanning (amnesic group mean: 14.9 days±0.86 amnesic group range: 13-28 days; control group mean: 16.1 days±3.21, range: 13-62; U=157, p=0.888). Episodic memories were sampled from a mean period of 5.28±0.51 years before symptom onset (range: 2011-2020) in the group with hippocampal amnesia. The control group were matched to this premorbid time period (mean period of 5.50±0.42 years before the amnesic group symptoms onset; range: 2012-2020). No between-group differences were found in the absolute age of these memories (amnesic group median: 5 years; controls median: 5; U=177, p=0.630). All participants were instructed to select three memories that were orthogonal to each other in terms of the content related to the people involved, the places, and the events to ensure they did not include any overlapping mnemonic details. All memories were checked at the point of acquisition for this degree of orthogonalisation for all participants, with no memories being rejected on the basis of these criteria. Episodic and semantic details associated with each of the three autobiographical memories were assessed under the retrieval conditions of the Autobiographical Interview procedure.^56^ Unique cue words were then agreed between experimenter and participant to serve as a basis for retrieval cues that were to be presented during the fMRI scan section of the experiment.

### Scoring and reliability

All verbal responses on the AI procedure were recorded in a digital audio format and then transcribed for scoring offline. Transcripts were compiled so that identifying personal details or anything that pertained to group membership were removed. Responses were scored according to the standardised method outlined in the AI Scoring Manual.^56^ Accounts were segmented into informational bits/details, or those occurrences, observations, or thoughts expressed as a grammatical clause. Details that related directly to a unique event, and which had a specific time and place or were associated with episodic re-experiencing (such as thoughts or emotions), were classified as internal (episodic) details. Information that did not relate to the event was assigned to external details, and then sub-categorized into semantic (factual information or extended events), repetitions (where previous details had been given with no new elaboration), and other (e.g., metacognitive statements, editorializing, and inferences).

One rater scored 100% of the episodes acquired with another rater scoring 50% of the acquired episodes (including post-scan debrief; see below), which is in line with prior studies.^52,56^ Intra-class correlation coefficients (ICCs) were used to calculate the agreement between the twice-repeated scoring of data acquired on the AI. A two-way mixed-model design was used to test the degree of absolute agreement, with the intraclass correlation coefficient of 0.94 indicated a high-level of consistency in scoring.

### Autobiographical Interview statistical analysis

In line with previous work,^31,37^ we analysed the internal (episodic) and external (semantic) points generated per participant with an omnibus ANOVA and followed up with planned comparisons. The alpha criterion was corrected for multiple comparisons using the Bonferroni-Holm procedure.

### Task and procedure

The neuroimaging section of the experiment consisted of four parts: a pre-scan training session; an intra-scan training session; the autobiographical retrieval session; and a post-scan debriefing session. Experimental tasks were run on a Dell 15-inch laptop using PsychoPy software (v.2022.2.4.),^169^, with stimulus presentation delivered via a MRI-compatible projection system (JVC DLA-SX21 projector via HDMI cables; 1400×1050 pixels resolution run at native resolution and a native refresh rate of 60Hz). All task stimuli were viewed on a 26×19.5cm screen positioned at a distance of 62cm from the participant. Tones were played through the Etymotic Ear-Tone stereo sound system.

### Pre-scan training

All participants underwent training in a behavioural testing room located outside of the MRI scan room. Each pre-scan training session comprised three runs. Training runs were based on two other memories acquired during the autobiographical memory acquisition phase conducted two-weeks prior to the scan session. On the first training run, participants were presented with the title of a retrieval cue for a specific memory. Each cue appeared on a computer screen for three seconds and was followed by the presentation of an on-screen instruction to ‘Close your eyes’ (12 seconds). Participants were instructed to close their eyes as soon as they knew which memory they had to remember and then verbally recount the memory with as much vivid detail as possible until a tone sounded. The prompt to remember and recount each episodic memory was repeated three times and in a pseudorandom order, such that each retrieval cue prompt was not presented twice in succession. The participants then had a rest screen for five-seconds before the next trial. The second training run comprised the same three memories, with each remembered three times out loud over the 15s total retrieval period. On the second run, after recounting the memory, participants were additionally asked to indicate the vividness associated with each specific memory on a four-point scale (1 – very vivid, as if like real life, 2 – vivid but not like real life, 3 – some recollection, and 4 – no meaningful experience of the memory).

Participants were asked to provide the rating within three-second interval. After each rating, participants were then provided with a five-second rest period before the next trial. The third training run consisted of the same three practice memories with vividness ratings, each retrieved verbally three times, with the addition of a control task that was psuedorandomly interleaved with the remember trials. On the control task, participants were instructed to count up from a given starting number in a stated interval and continue counting for 15s. After each control trial, the participants were asked to rate their level of task engagement on a four-point scale (1 – entirely engaged with counting, 2 – mostly engaged with counting, 3 – some counting, and 4 – completely distracted).

During the final training run, participants were first provided with a reminder about their chosen memories and the titles associated with each memory. During this training block, participants were presented with the titles as retrieval cues for their experimental memories. All participants were encouraged to begin memory retrieval during that cue screen as detailed above, except this time all participants were asked to retrieve their memories with as much visual detail as possible and to remember silently. All the participants were able to retrieve their specific memories. Total retrieval time was once again 15s for each memory. These memories were psuedorandomly interleaved with the counting task. Both trial types were followed by their respective rating assessment and then a five-second rest period.

### Intra-scan training

All participants underwent a final training session on the autobiographical memory remembering task once insider the MRI scanner. During this final training phase, participants were asked to remember each autobiographical memory in silence with their eyes closed for the three recollections in order to align with the upcoming experimental conditions. This was used to familiarise all participants to the conditions of fMRI phase of the experiment. At the end of each trial, participants were asked to provide ratings for the memory and counting trials, as detailed above, and input their numerical response via an MRI-compatible button box.

### Autobiographical memory task

During the acquisition of fMRI data, participants underwent three trial runs in which each memory retrieval cue was presented five-times per block, interleaved with the counting task. The presentation of each stimulus was pseudo-randomised such that each condition was not presented in succession. In total, there were 15 trials for each memory, totalling 45 memory trials performed under fMRI conditions.

### Post-scan debriefing session

At the end of the fMRI scan session, each participant underwent another recorded Autobiographical Interview procedure, where they were asked to remember what they were visualising for each memory, in as much detail as possible, on each of the memory trials. The debriefing sessions were transcribed and scored in the same manner as the Autobiographical Interview.

### Functional and anatomical MRI data acquisition

Structural and functional MRI data were acquired using a Siemens 3.0-Tesla Prisma MRI system (Magnetom TIM Trio, Siemens Healthcare, Erlangen, Germany) with a 64-channel head coil.

Functional MRI data were collected with a T2*-weighted echo-planar imaging (EPI) sequence (TR = 3.36s, TE = 30 ms, slice thickness = 2.5 mm, 3 x 3mm in-plane resolution, phase encoding direction = anterior → posterior, field of view = 192 mm^2^, matrix size = 64 x 72, flip angle = 90°, slice tilt = -30°, 48 axial slices), resulting in total of 210 whole-brain volumes per run (lasting for ∼12 minutes per run).

A whole-brain structural T1-weighted sequence was acquired (six-minute acquisition time) after the experimental EPI runs for each participant at an isotropic resolution of 0.8mm,^170^ which was used for the automated VBM analysis and FreeSurfer cortical parcellation. Whole-brain T1-weighted anatomical images consisted of 176 axial slices acquired with a magnetisation-prepared rapid gradient-echo (MPRAGE) sequence (TI = 1100 ms, TR = 2530 ms, TE = 3.34 ms, slice thickness = 1 mm, 1 mm^2^ in-plane resolution, phase encoding direction = anterior → posterior, field of view = 256 mm^2^, matrix size = 256 x 256, flip angle = 7°).

In order to conduct the semi-automated segmentation of the hippocampal subfields, partial volume images for the entire extent of the temporal lobes were collected using a single-slab 3D T2-weighted turbo spin-echo sequence with variable flip angles (SPACE)^171^ in combination with parallel imaging, to simultaneously achieve a high image resolution of ∼500 μm, high sampling efficiency, and short scan time while maintaining a sufficient (SNR). After excitation of a single axial slab, the image was read out with the following parameters: resolution = 0.5 × 0.5 × 0.5 mm, matrix = 384 × 328, partitions = 104, partition thickness = 0.5 mm, partition oversampling = 15.4%, field of view = 200 × 171 mm^2^, TE = 353 ms, TR = 3200 ms, GRAPPA x 2 in phase-encoding (PE) direction, bandwidth = 434 Hz/pixel, echo spacing = 4.98 ms, turbo factor in PE direction = 177, echo train duration = 881, averages = 1.9, plane of acquisition = sagittal. To improve the SNR of the anatomical image, two scans were acquired for each participant (taking 13m each), coregistered and averaged. For each participant, the 3D T2-weighted scans underwent Rician noise estimation^172^ and were then denoised using oracle-based discrete cosine transform (ODCT^173^), with additional denoising then applied to the ODCT denoised image using a prefiltered rotationally invariant nonlocal means filter.^173^ This computed a single denoised image for each high-resolution temporal lobe image. The denoised images were then co-registered and averaged to provide a final image for hippocampal segmentation for each participant (see below).

### Whole-brain voxel-by-voxel morphometry and cortical parcellation

Whole-brain voxel-by-voxel morphometry (VBM) and diffeomorphic anatomical registration using the exponentiated Lie algebra (DARTEL) registration method^174^ were conducted on the 3.0-Tesla (0.8 x 0.8 x 0.8mm^3^ spatial resolution) T1-weighted anatomical images, using grey matter volume, white matter volume, and cerebrospinal fluid volumes as covariates. The voxel-by-voxel contrast of normalized grey matter across the whole-brain was conducted using a two-sample *t*-test thresholded at p<0.05, with family-wise error correction for multiple comparisons. The results revealed no suprathreshold clusters of grey matter or white matter volume loss, nor any increase in cerebrospinal fluid volume, in the group with hippocampal amnesia relative to the control group.

T1-wighted images were also segmented and parcellated into cortical and subcortical regions of interest (ROI) based on the Destrieux Atlas^175^ using the -recon-all function in FreeSurfer version 7.3.2 (http://surfer.nmr.mgh.harvard.edu/). This atlas was chosen as it contains a greater number of cortical parcellations that correspond to the specific cytoarchitectural boundaries constituting Brodmann areas.^175^ In this way, we limited the spatial extent of voxels within a given region-of-interest (ROI), facilitating a more fine-grained analysis.^176^ Following cortical parcellation, each T1-weighted image obtained for each participant and the respective parcellations were transformed into the participant’s native anatomical space using a rigid boundary-based registration approach, specific for registration between EPI and structural images.^177^

### Anatomical segmentation of hippocampal subfields

3D T2-weighted turbo spin-echo images provided the basis for automated segmentation using the Automatic Segmentation of Hippocampal Subfields (ASHS^178^) atlas package.^179^ The ASHS atlas training process works by first applying deformable coregistration of the T1-weighted image, T2-weighted image and hippocampal subfield masks to an unbiased population template. Automatic segmentations are then produced by deformably coregistering the T2-weighted image of each participant to that of all other participants, and applying joint label fusion.^180^ Finally, corrective learning classifiers^181^ are trained by comparing the automatic segmentations with the manual segmentation of the same T2-weighted image; see Yushkevich et al. 2015^178^ for full details of the ASHS pipeline. Once participant scans had been processed through the ASHS pipeline, all resultant hippocampal segmented images were then manually inspected and modified according to a previously described methodology^182^ using the ITK Snap software (version 3.8.0). Masks were created for the following six subfields: DG/CA4, CA2/3, CA1, subiculum, pre/parasubiculum and uncus. Subfield segmentations were performed by an experienced researcher (TDM). Volumes were then corrected for intracranial volume in line with our previous work.^31,37^ To evaluate intra-rater reliability, five datasets from participants in the group with hippocampal amnesia and five datasets from participants in the control group (>10% of experimental population) were re-segmented after a month had elapsed. Analyses for each subfield were conducted using the Dice overlap metric.^183^ Dice scores are reported in Supplementary Table 1, and ranged from 0.7259-0.9391. Intraclass correlation coefficients were derived from the 12 original and replicated subfield volumes across both segmentations (24 volumes per participant) and across both groups (240 subfield volumes in total) using a one-way random effects model.

### Hippocampal subfield segmentation statistical analysis

In line with previous work,^31,37^ hippocampal subfield volumes were examined using a mixed-model omnibus ANOVA and planned between-group comparisons

### Autobiographical network definition

Given the overlap between the default mode network, the autobiographical memory retrieval network,^91,92,102,125,145,184–186^ and prior evidence of altered graph theoretical metrics within these regions in a cohort of individuals with amnesia secondary to LGI-LE,^31^ we sought to evaluate the RDM model fit and trial-by-trial similarity in the discrete pattern of voxels that might correspond to specific autobiographical memories within this network. Moreover, there is increasing evidence that informational components of episodic memories are supported by discrete networks, such as information about people within the anterior temporal network,^14^ contextual information in the posterior medial network,^14^ schema in the medial prefrontal network,^14^ and event-specific representations within the hippocampus.^14^ Therefore, in order to examine representational similarity within these regions, we also extended the cortical network to structures of the dorsolateral PFC (dlPFC), because the dlPFC plays a critical role within autobiographical memory retrieval,^92,109,111,120,187,188^ and also appears to constitute an extended component of the default mode network.^189^ Therefore, representational similarity for autobiographical memories was examined in the following regions (Destrieux Atlas labels are provided within parentheses): anterior medial prefrontal cortex (G_Orbital); posterior cingulate cortex (G_Cingul_Postdorsal); temporoparietal junction (G_occipital_middle); angular gyrus (G_parietalAngular); orbitofrontal cortex (medial and lateral; G_orbital), inferior frontal gyrus (G_Front_inf-Orbital), precuneus (G_precuneus), retrosplenial cortex (S_pericallosal), middle frontal gyrus (G_front_middle), lateral temporal cortex (G_temporal_middle), ventromedial prefrontal cortex (G_rectus), parahippocampal cortex (G_oc-temp_med-Parahip). All regions were evaluated in both hemispheres on the basis that left and right hemispheric structures have distinct roles in AM and episodic memory retrieval.^92,101,102,109,113,120,190^

### Functional magnetic resonance pre-processing

MRI data were pre-processed using SPM12 (http://fil.ion.ucl.ac.uk/spm/). The first 5 volumes of each functional run were discarded to allow for T1 equilibration. Functional images were unwarped using the collected field maps^191^ and slice-time corrected but left unsmoothed and in native space. Structural T1-weighted images were coregistered to the mean functional image of each participant. Each participant’s structural image was segmented into grey matter, white matter, and cerebrospinal fluid using a nonlinear deformation field to map it onto a template tissue probability map.^192^ Motion correction parameters estimated from the re-alignment procedure and their first temporal derivatives (12 regressors in total) were included as confounds in the first-level analysis for each participant.

### Representational similarity analysis

Representation Similarity Analysis (RSA) is a method used in multivariate pattern analysis (MVPA) to examine how similar or dissimilar the activity patterns are for the same versus different stimuli.^61,62^ For each pair of trials, the multivariate distance (or similarity) between their activity patterns was measured using Pearson correlation. High similarity (high correlation or low distance; also denoted as low dissimilarity) indicates that the multivoxel patterns for the two trials are similar and may therefore involve the same set of neurons. Therefore, RSA provides an indirect measure of representational content within neuronal populations that is then associated with cognition. Unlike the standard mass-univariate analysis, which examines each voxel independently, RSA considers the joint activity of multiple voxels to decode information about experimental conditions. This allows for the detection of groups of voxels that represent very specific and subtle experimental conditions, such as specific episodic memories. There are several reasons for the high sensitivity afforded by MVPA: (1) RSA can combine weak but consistent signals from multiple voxels; distributed networks of very weakly active voxels that show no univariate effect may be detectable by RSA; and (2) it can measure the relationship between distal voxels where the relationship in their activity is required for memory representation, but the BOLD signal change might not be significant on univariate analysis.^14,42^

All RSAs were run on unsmoothed native-space functional images after the preprocessing steps described above had been completed. Each of the three retrieval runs were entered into a general linear model (GLM) in SPM to directly mitigate influences of signal drift, motion, and global nuisance signal in evaluating representational patterns. Each memory retrieval event was modelled separately from all other retrieval events both within runs and across runs. As memory retrieval events, including title screens, were 15s in duration and the TR for our EPI sequence was 3.36s, there were approximately four TRs per retrieval, but 67 TRs per memory condition (201 TRs for the episodic retrieval and control task conditions). The beta values were divided by the square root of the GLM residuals to reduce the impact of noisier voxels._193_ Resultant beta images were masked using a region-of-interest (ROI) approach, based on the network definition listed above. Voxels with null values in any scan run were excluded. A split representational dissimilarity approach ^14,194^ was undertaken, with analyses conducted so that we computed retrieval-by-retrieval pattern similarity for each pair of trials (memory and counting, using trial-specific beta values) within runs and then these were averaged across the three runs.

### Model matrix comparisons

We aimed to compare memory-by-memory correlations using two specific model hypothesis matrices to assess the following: (1) whether the observed hippocampal subfield pathology resulted in between-group differences in representational content during episodic memory retrieval, as a general process compared to the control task, across the network of ROIs (see Figure 4A); and, (2) whether hippocampal subfield pathology generated between-group differences in the capacity to represent specific episodic memories across the network of ROIs (see Figure 4B). Each run comprised 30 trials (15 trials per condition, with five trials per specific memory). Therefore, RDM model fit correlation scores were created from 465 within-run trial-by-trial correlation pairs (comparing each condition to all of the other trials per run), then averaged across runs. For the specific episodic memory retrieval hypothesis matrix, the counting trials were also modelled within the hypothesis matrix (creating a 30 x 30 matrix of trial-by-trial correlations), with the diagonal representing low dissimilarity. This is not shown in Figure 4B for simplicity. Model matrices were constructed to feature a value of 1 in hypothesized high-dissimilarity cells and a value of 0 in hypothesized low-dissimilarity cells. Beta images resulting from the preprocessing steps above, characterising the amplitude of the BOLD response to each trial, underwent pairwise comparisons to yield a Pearson Z dissimilarity score (computed as 1-Pearson correlation score) for each trial-by-trial comparison. Model fits were evaluated using a correlation between the pattern similarity matrix and the model matrices for each ROI using Kendall τ (the memories were modelled as categorical data as well as being a smaller sample set; tested using a stimulus-label randomization test by randomly permuting stimulus order for each subject 10,000 times to create a distribution of permuted within-subjects correlations under the null hypothesis that the hypothesis matrices could not be decoded, thresholded to a false detection rate of p<0.05). Model fits were then used in two separate ways: (1) to evaluate whether, for each group, the group level model fit was significantly different from the random condition using permutation testing, thereby suggesting the hypothesis matrix correctly modelled the multivoxel-patterned data; and, (2) as the basis to assess whether there was a between-group difference in degree of model fit, whereby we hypothesised that participants in the group with hippocampal amnesia would have significantly lower model fits within our ROIs. To our knowledge, there is no robust evidence to suggest that correlation values within different brain regions for the same task should be related, so between-group differences were assessed using between-group *t*-tests and orthogonality was assumed between the ROIs. Where a significant between-group difference was found for the specific episodic memory hypothesis matrix, we subjected the first-level Fisher-transformed Pearson scores for the individual memories (averaging the Fisher transformed Pearson correlation scores for each pair of within-memory trials for the same within individual runs, and then across runs) to a 2 (group: amnesic, controls) x 3 (experimental memories probed: 1, 2, and 3) omnibus ANOVA to assess whether any differences in model fit were represented in the first-level scores for that analysis.

We also sought to understand the relationship between these RSA results with the volumetric hippocampal subfield results and the internal (episodic) memory detail scores as detailed above using linear regression modelling, in line with our previous work.^31,37^

### Statistical analyses

All t-tests, ANOVAs, and linear regressions were performed on IBM SPSS Version 29.0 (IBM Corp.)

## Acknowledgements

This work is dedicated to Professor Eleanor Maguire, who was involved throughout the experiment, demonstrating her courageousness and dedication to cognitive neuroscience. Eleanor passed away before manuscript completion. The work reported here represents one of Eleanor’s final empirical contributions to the neuroscience of the hippocampus and episodic memory. Eleanor’s legacy will be continued by her trainees and collaborators. We would like to thank Professor Cathy Price for her comments on the manuscript.

TDM is supported by the Wellcome Trust (Grant number: 222913/Z/21/Z). AEH is supported by the Medical Research Council (MR/X022013/1 and MR/V007173/1), and the National Institute for Health Research (NIHR) Oxford Health Biomedical Research Centre (BRC).

## Author contributions

TDM: conceptualisation, methodology, investigation, data curation, formal analysis, validation. funding acquisition, resources, project administration, visualisation, and writing – original draft, reviewing and editing; ALH: methodology, software, formal analysis, and writing – reviewing and editing; YIW: methodology, software, and writing – reviewing and editing; JZ: formal analysis and writing – reviewing and editing; AEH: resources, writing – reviewing and editing; EC: resources, writing – reviewing and editing; TAP: resources, writing – reviewing and editing; MSZ: resources and writing – reviewing and editing; CRR: formal analysis, visualisation, writing – original draft, reviewing and editing; EAM: conceptualisation, methodology, formal analysis, funding acquisition, resources, project administration, and supervision.

For the purpose of Open Access, the author has applied a CC BY public copyright licence to any Author Accepted Manuscript version arising from this submission.

## Additional information Supplementary information

**Correspondence** and requests for materials should be addressed to Thomas D. Miller.

