## Supplementary data for "CA2/3-dependent stability of frontoparietal mnemonic representations predicts episodic deficits in human amnesia"

### Supplementary information

#### Results

#### Neuropsychological assessment

All participants underwent extensive standardised neuropsychological assessment in line with our previous work (detailed in the Methods section). Supplementary Table 1 details the performance of the group with hippocampal amnesia compared to the normative *Z*-transformed data set (with a mean of 0 and standard deviation of 0.3, tested with a between-group independent sample two-tailed *t*-test). Supplementary Table 2 details the neuropsychological domain *Z*-score performance between the hippocampal amnesia and control groups (assessed with between-group independent-sample two-tailed *t*-test). One participant in the group with hippocampal amnesia could not undertake the Test of Premorbid Functioning (TOPF) due to severe dyslexia.

Briefly, the results demonstrated that, compared to the normative data (Supplementary Table 1), the group with hippocampal amnesia exhibited significantly superior performance for verbal and visual intelligence, premorbid intelligence, language, executive function, cognitive flexibility, and visuomotor skills. Only visual memory was significantly impaired in the group with hippocampal amnesia.

**Supplementary Table 1. Neuropsychological domain performance of the LGI1-LE (amnesic) group compared to normative data**

| Domain | n | Average Z-score | SEM | *t* | d.f. | *p*-value |
|---|---|---|---|---|---|---|
| Verbal intelligence | 18 | 1.04 | 0.08 | 4.4 | 17 | <0.001 |
| Visual intelligence | 18 | 0.87 | 0.15 | 3.69 | 17 | <0.001 |
| Premorbid intelligence | 17 | 0.84 | 0.15 | 3.48 | 16 | <0.001 |
| Verbal memory | 18 | 0.26 | 0.27 | 1.1 | 17 | 0.35 |
| Visual memory | 18 | **-0.92** | **0.16** | **-3.91** | **17** | **<0.001** |
| Recognition memory | 18 | 0.13 | 0.27 | 0.55 | 17 | 0.64 |
| Sustained attention | 18 | 0.28 | 0.22 | 1.18 | 17 | 0.22 |
| Language | 18 | 0.89 | 0.26 | 3.77 | 17 | 0.003 |
| Executive function | 18 | 0.91 | 0.17 | 3.85 | 17 | <0.001 |
| Cognitive flexibility | 18 | 0.81 | 0.17 | 3.46 | 17 | <0.001 |
| Visuomotor skills | 18 | 0.35 | 0.14 | 1.49 | 17 | 0.02 |
| Visuoconstruction | 18 | 0.32 | 0.18 | 1.36 | 17 | 0.087 |

Bold results indicate group-level neuropsychological domains significantly impaired in the amnesic group compared to the normative control population means.

When compared to the control cohort in this study (Supplementary Table 2), the group with hippocampal amnesia exhibited significant impairment in verbal memory (patients' mean: 0.26±0.27; controls' mean 1.54±0.24; $t$ = -3.56, $p$<0.001, Cohen's d = -1.19) and in visual memory (patients' mean: -0.92±0.16; controls' mean: -0.20±0.13); $t$ = -3.52, $p$<0.001, Cohen's d = -1.17). These results indicate that there were statistically significant differences between the hippocampal amnesia and control groups, which were associated with large effect sizes.

**Supplementary Table 2. Neuropsychological domain performance of the LGI1-LE amnesic group compared to the control group**

| **Domain** | **n** | **Amnesic group average Z-score (SEM)** | **Control group average Z-score (SEM)** | ***t*** | **d.f.** | ***p*-value** |
|---|---|---|---|---|---|---|
| Verbal intelligence | 18 | 1.04 (0.08) | 1.32 (0.13) | -1.84 | 34 | 0.074 |
| Visual intelligence | 18 | 0.87 (0.15) | 1.26 (0.20) | -1.54 | 34 | 0.13 |
| Premorbid intelligence | 17 | 0.84 (0.15) | 1.10 (0.11) | -1.42 | 33 | 0.17 |
| Verbal memory | 18 | **0.26 (0.27)** | **1.54 (0.24)** | **3.56** | **34** | **<0.001** |
| Visual memory | 18 | **0.92 (0.16)** | **0.20 (0.13)** | **-3.52** | **34** | **<0.001** |
| Recognition memory | 18 | 0.13 (0.27) | 0.50 (0.18) | -1.14 | 34 | 0.26 |
| Sustained attention | 18 | 0.28 (0.22) | 0.04 (0.12) | 0.97 | 34 | 0.34 |
| Language | 18 | 0.89 (0.26) | 1.43 (0.22) | -1.61 | 34 | 0.12 |
| Executive function | 18 | 0.73 (0.17) | 1.00 (0.18) | -1.08 | 34 | 0.29 |
| Cognitive flexibility | 18 | 0.82 (0.17) | 0.85 (0.15) | -0.17 | 34 | 0.87 |
| Visuomotor skills | 18 | 0.35 (0.14) | 0.52 (0.13) | -0.87 | 34 | 0.39 |
| Visuoconstruction | 18 | 0.32 (0.18) | 0.59 (0.15) | -1.19 | 34 | 0.24 |

Bold results indicate group-level neuropsychological domains significantly impaired in the amnesic group compared to the control group.

**Selective bilateral hippocampal subfield volume loss**

Supplementary Table 3 reports the intraclass correlation coefficients and Dice scores for the semi-automated segmentation of the hippocampal subfields. Supplementary Table 4 reports the specific group level bilateral subfield volumes from the amnesic and control participants groups after semi-automated hippocampal subfield segmentations. Mean total subfield volumes are reported with standard error of the mean and standard deviation alongside the results from the planned comparisons conducted to assess between-group difference in hippocampal subfield volumes. Mean total subfield volumes were collapsed across left and right hemispheres because was no significant interaction between group, side (left, right), and subfield [$F_{(3.679,125.082)} = 0.894$, $p = 0.486$, $\eta^2 = 0.038$).

**Reliability of hippocampal subfield segmentations**

**Intraclass correlation coefficients for the Dice scores** (see Supplementary Table 3) indicated a high degree of reliability in the output of two repetitions of the segmentation protocol in a subset (n = five per group) of participants from the group with hippocampal amnesia and from the control group. Intraclass correlation coefficients were derived from the 12 original and replicated subfield volumes per participants (24 subfield volumes in total) and across the 10 re-segmented participants (creating 240 subfield volumes in total) using a one-way random effects model. This demonstrated an average measures intraclass correlation across both groups of 0.813 (controls: 0.942; amnesic group: 0.905).

**Supplementary table 3. Dice scores for semi-automated manual hippocampal subfield segmentation**

| **Group** | **Side** | **Subfield** | **Average (median; range)** |
|---|---|---|---|
| Controls | Left | Dentate gyrus | 0.864±0.025 (0.854; 0.7898-0.9363) |
| | | CA1 | 0.900±0.012 (0.903; 0.864-0.936) |
| | | CA23 | 0.821±0.023 (0.845; 0.7610-0.8730) |
| | | Subiculum | 0.872±0.021 (0.876; 0.8176-0.9276) |
| | | Pre/parasubiculum | 0.846±0.031 (0.868; 0.7267-0.8956) |
| | | Uncus | 0.897±0.024 (0.896; 0.8171-0.9703) |
| | Right | Dentate gyrus | 0.882±0.034 (0.877; 0.8717-0.9551) |
| | | CA1 | 0.898±0.014 (0.894; 0.8650-0.9347) |
| | | CA23 | 0.856±0.032 (0.867; 0.7537-0.9347) |
| | | Subiculum | 0.900±0.021 (0.916; 0.8256-0.9506) |
| | | Pre/parasubiculum | 0.851±0.018 (0.858; 0.8012-0.8883 |
| | | Uncus | 0.888±0.019 (0.891; 0.8266-0.9391) |
| Amnesic group | Left | Dentate gyrus | 0.895±0.008 (0.887; 0.8737-0.9188) |
| | | CA1 | 0.895±0.010 (0.899; 0.8623-0.9186) |
| | | CA23 | 0.836±0.011 (0.841; 0.8009-0.8712) |
| | | Subiculum | 0.885±0.015 (0.897; 0.8357-0.917) |
| | | Pre/parasubiculum | 0.847±0.009 (0.845; 0.8285-0.8771) |
| | | Uncus | 0.868±0.018 (0.874; 0.8055-0.9145) |
| | Right | Dentate gyrus | 0.876±0.012 (0.866; 0.8452-0.9159) |
| | | CA1 | 0.885±0.014 (0.891; 0.8476-0.8911) |
| | | CA23 | 0.810±0.022 (0.819; 0.7259-0.8547) |
| | | Subiculum | 0.895±0.011 (0.892; 0.8661-0.9215) |
| | | Pre/parasubiculum | 0.819±0.018 (0.834; 0.7557-0.8574) |
| | | Uncus | 0.814±0.017 (0.808; 0.7638-0.8720) |

**Supplementary Table 4. Hippocampal subfield volumes in the group with hippocampal amnesia (LGI1-LE patients) group and controls (n=18 in both groups).** Volumes were normalised to the total intracranial volumes obtained from the voxel-based morphometry analyses. Volumes collapsed across both sides as there was no significant interaction between group, side (left, right), and subfield.

**Supplementary table 4. Hippocampal subfields volumes in the amnesic (LGI1-LE) group and control group (n = 18)**

| | **Mean total subfield volumes** | | |
|---|---|---|---|
| **Subfield** | **Amnesic group** | **Control group** | **Planned comparison (% difference)** |
| CA1 | 729.79 (25.77, 109.33) | 891.46 (30.76, 130.50) | Significant (-18.13) |
| CA23 | 156.46 (4.32, 18.34) | 263.07 (8.01, 33.97) | Significant (-40.53) |
| Dentate gyrus | 637.63 (25.26, 107.18) | 673.94 (28.13, 119.34) | ns (-5.39) |
| SUB | 543.63 (26.65, 113.07) | 630.5 (21.63, 91.78) | ns (-13.73) |
| Pre/para SUB | 206.71 (8.31, 35.28) | 235.15 (5.89, 25.00) | ns (-8.81) |
| Uncus | 398.61 (25.46, 108.03) | 437.13 (16.49, 69.98) | ns (-12.10) |

Values are mean, $mm^3$ (±SEM, SD). Significant at Bonferroni correction for multiple comparisons following mixed-model ANOVA; ns = non-significant at alpha criterion corrected for multiple comparisons. CA1: cornu Ammonis 1; CA23: cornu Ammonis 2 & 3; ns: not significant; SUB: subiculum

**Intra-fMRI scan behavioural measures of visualisation (memory trials) and task engagement (counting trials)**

The subjective ratings given by participants after each memory ('how vivid was the recollection?') and the counting trials ('how engaged with the task were you') did not vary between the group with hippocampal amnesia and control group [memory rating: controls: 1.64±0.17; amnesic group 1.51±0.17; counting rating: controls: 1.60±0.16; amnesic group: 1.53±0.16]. Mauchly's test of sphericity indicated that the assumption of sphericity had been violated for trial type (memory versus counting) by run number (1-3) [$\chi^2_{(2)} = 9.480$, $p = 0.009$]. Therefore, the degrees of freedom were corrected using Huynh-Feldt estimates ($\varepsilon = 0.858$). There was no main effect of group ($F_{(1,34)} = 0.804$, $p = 0.376$, $\eta^2 = 0.023$), trial type ($F_{(1,34)} = 0.024$, $p = 0.878$, $\eta^2=0.001$), or run ($F_{(2,68)} = 2.367$, $p = 0.101$, $\eta^2 = 0.065$). There were no significant interactions between group and trial type ($F_{(1,34)} = 0.162$, $p = 0.690$, $\eta^2 = 0.005$), group and run ($F_{(2,68)} = 2.148$, $p = 0.125$, $\eta^2 = 0.059$), and trial type and run ($F_{(1.717,58.361)} = 1.889$, $p = 0.304$, $\eta^2 = 0.053$). The three-way interaction was also not significant ($F_{(1.717,58.361)} = 1.187$, $p = 0.310$, $\eta^2 = 0.034$).

**Post-scan event memory performance**

Following scanning, all participants underwent post-scan debriefing. This was undertaken in the form of the AI procedure, in which participants were asked to recall, in as much detail as possible, the content of their recollections during the fMRI scan. This was conducted with a general probe followed by a specific probe, as per the AI procedure handbook [internal (episodic) details: controls: 57.88±3.92; amnesic group 43.33±3.33; external (semantic) details: controls: 0.94±0.44; amnesic group: 0.61±0.33].

An omnibus 2 (group: amnesic, control) x 3 (memory: experimental event memories) x 2 [(memory detail type: internal (episodic), external (semantic)] mixed-model factorial ANOVA was conducted on the units of information acquired from three experimental memories, scored in accordance with AI procedure handbook after scanning. One control was excluded from this analysis as their accumulated internal detail score (7) was more than two standard deviations (within-group: 19.75; entire experimental population: 17.78) below the control (55.06) and entire experimental population mean (49.19). Mauchly's test of sphericity indicated that the assumption of sphericity had not been violated. There were significant main effects of group ($F_{(1,34)} = 8.936$, $p = 0.005$, $\eta^2 = 0.213$), memory ($F_{(2,66)} = 6.351$, $p = 0.003$, $\eta^2 = 0.161$), and memory detail type ($F_{(1,34)} = 369.845$, $p<0.001$, $\eta^2 = 0.918$). Group by memory detail type was significant ($F_{(1,34)} = 7.475$, $p = 0.010$, $\eta^2 = 0.185$). Planned comparisons demonstrated a significant difference in total internal (episodic) details ($F_{(1,34)} = 8.269$, $p = 0.007$, $\eta^2 = 0.200$) but not external (semantic) details ($F_{(1,34)} = 0.506$, $p = 0.482$, $\eta^2 = 0.015$).

### Representational similarity analysis model fits

Alterations in the trial-by-trial similarity of neuronal populations supporting representational content for remembered personal events were assessed using representational dissimilarity matrices. The rationale for this analysis was two-fold: (1) to assess whether the patterned activity during episodic memory retrieval within our regions-of-interest (ROIs) were comparable to our hypothesised model matrices that depicted the two analytical conditions for the experiment (episodic memory retrieval as a general process and the retrieval of specific episodic memories); and, (2) to then assess where those representations within the defined AN exhibited between group differences. The first result reported in detail here is that both groups had significant model fits for both hypothesis matrices across our defined network. The between-group differences for both hypotheses are reported in the main text. Here, we find that the significant model fits with no between-group differences. All results are derived from between-group independent sample two-tailed *t*-tests on the participants' derived Kendall τ scores.

#### 1. Model fit: episodic memory retrieval compared to the control (counting) condition

#### Anterior medial prefrontal cortex (amPFC)

In the left anterior medial frontal cortex, the group with hippocampal amnesia showed a mean Kendall τ = 0.277±0.017 and the control group had a mean Kendall τ = 0.290±0.017. The between-group comparison yielded a non-significant difference ($t_{(34)} = 0.458$, $p = 0.650$, Cohen's d = 0.153). For the right amPFC, the group with hippocampal amnesia demonstrated a mean Kendall τ = 0.259±0.019, compared to the control group's mean Kendall τ = 0.285±0.020. The between-group contrast for the right amPFC was not significant ($t_{(34)} = 0.915$, $p = 0.366$, Cohen's d = 0.305). These results indicate that there were no statistically significant differences between the hippocampal amnesia and control groups in either the left or right amPFC, with small to moderate effect sizes observed.

#### Posterior cingulate cortex (PCC)

In the left PCC, the mean Kendall τ = 0.398±0.016 in the group with hippocampal amnesia compared to the control group's mean Kendall τ = 0.411±0.017. The between-group contrast for the left PCC was not significant ($t_{(34)} = 0.547$, $p = 0.588$, Cohen's d = 0.182). In the right PCC, the mean in the group with hippocampal amnesia was Kendall τ=0.373±0.017, whereas the mean Kendall τ for the control group was 0.396±0.018. Comparison between groups for the right PCC revealed a non-significant difference ($t_{(34)} = 0.927$, $p = 0.360$, Cohen's d = 0.309). These findings suggest that there were no statistically significant differences between the hippocampal amnesia  and control groups in either the left or right PCC, with effect sizes that were small to moderate.

**Temporoparietal junction (TpJ)**

In the left TpJ, the mean Kendall τ = 0.268±0.025 in the group with hippocampal amnesia compared to the control group's mean Kendall τ = 0.320±0.021. The between-group contrast for the left temporoparietal junction was not significant ($t_{(34)}$ = 1.048, $p$ = 0.302, Cohen's d = 0.349). In the right TpJ, the mean Kendall τ = 0.307±0.025 in the group with hippocampal amnesia in contrast to the control group's mean Kendall τ = 0.327±0.016. The comparison between groups for the right TpJ was not significant ($t_{(34)}$ = 0.662, $p$ = 0.512, and Cohen's d = 0.221). These findings indicate no statistically significant differences between the group with hippocampal amnesia and control groups in either the left or right TpJ, with small to moderate effect sizes observed.

**Right angular gyrus**

In the right angular gyrus, the mean Kendall τ=0.375±0.019 in the group with hippocampal amnesia, whereas the mean Kendall τ:=0.422±0.015 for the control group. The between-group contrast for the right angular gyrus was not significant ($t_{(34)}$ = 1.883; $p$ = 0.068, Cohen's d = 0.628). These results indicated that there were no statistically significant differences between the hippocampal amnesia and control groups in right angular gyrus, with small to moderate effect sizes observed.

**Orbitofrontal cortex (medial; mPFC)**

In the left orbitofrontal cortex, the mean Kendall τ=0.277±0.096 in the group with hippocampal amnesia, whereas the control group had a mean Kendall τ=0.290±0.017, and there was no significant difference between the groups($t_{(34)}$ = 0.458, $p$ = 0.650, Cohen's d = 0.153). Similarly, in the right orbitofrontal region, the mean Kendall τ=0.259±0.019 in the group with hippocampal amnesia compared to the control group's mean Kendall τ=0.285±0.020. Again, there was no significant difference between the groups ($t_{(34)}$ = 0.915, $p$ = 0.366, Cohen's d = 0.305). These results indicate that there were no statistically significant differences between the hippocampal amnesia and control groups in either the left or right mPFC, with small to moderate effect sizes observed.

**Inferior frontal gyrus (IFG)**

The study also examined the left and right inferior frontal gyri in the amnesic group and control group. For the left inferior frontal gyrus, the mean Kendall τ = 0.284±0.021 in the group with hippocampal amnesia, compared to 0.330±0.021 in the control group. No significant difference was obtained between the groups ($t_{(34)}$ = 1.558, $p$ = 0.128, Cohen's d = 0.5198). Similarly, in the right inferior frontal gyrus, the mean Kendall τ = 0.270±0.022 in the group with hippocampal amnesia, compared to the mean Kendall τ = 0.314±0.022 in the control group. Once again, no significant between-group difference was found ($t_{(34)}$ = 1.388, $p$ = 0.174, Cohen's d = 0.463). These results indicate that there were no statistically significant differences between the hippocampal amnesia and control groups in either the left or right IFG, with small to moderate effect sizes observed.

**Precuneus**

In the left precuneus, the mean Kendall τ = 0.404±0.017 in the group with hippocampal amnesia, whereas the mean Kendall τ = 0.433±0.011 in the control group. The mean Kendall τ values were not significant between-groups ($t_{(34)}$ = 1.437, $p$ = 0.160, Cohen's d = 0.479). In the right precuneus, the mean Kendall τ = 0.371±0.020 in the group with hippocampal amnesia, compared to 0.406±0.015 in the control group. Once again, the difference between groups did not reach significance ($t_{(34)}$ = 1.405, $p$ = 0.169, Cohen's d = 0.468). These results indicate that there were no statistically significant differences between the hippocampal amnesia and control groups in either the left or right precuneus, with small to moderate effect sizes observed.

**Retrosplenial cortex**

In the left retrosplenial cortex, the mean Kendall τ = 0.305±0.017 in the group with hippocampal amnesia compared to 0.336±0.011 in the control group. The between-group difference was not significant ($t_{(34)}$ = 1.507, $p$ = 0.113, Cohen's d = 0.502). In the right retrosplenial cortex, the mean Kendall τ=0.277±0.017 in the group with hippocampal amnesia compared to 0.316±0.014 in the control group, which was not significantly different between groups ($t_{(34)}$ = 1.841, $p$ = 0.074, Cohen's d = 0.614). These results indicate that there were no statistically significant differences between the hippocampal amnesia and control groups in either the left or right retrosplenial cortex, with small to moderate effect sizes observed.

**Middle frontal gyrus (MFG)**

In the left middle frontal gyrus, the mean Kendall τ = 0.364±0.024 in the group with hippocampal amnesia compared to 0.421±0.014 in the control group, which was not significantly different between groups ($t_{(34)}$ = 2.001, $p$ = 0.053, Cohen's d = 0.667). In the right middle frontal gyrus, the amnesic group's mean Kendall $\tau$=0.317±0.021 compared to 0.368±0.020 in the control group. The between-group difference did not reach significance ($t_{(34)}$ = 1.714, $p$ = 0.096, Cohen's d = 0.571). These results indicate that there were no statistically significant differences between the hippocampal amnesia and control groups in either the left or right MFG, with small to moderate effect sizes observed.

**Lateral temporal cortex (LTC)**

In the left lateral temporal cortex, the mean Kendall $\tau$ = 0.341±0.023 in the group with hippocampal amnesia, whereas in the control group the mean Kendall $\tau$ = 0.381±0.021. The between-group difference was not significant ($t_{(34)}$ = 1.286, $p$ = 0.208, Cohen's d = 0.429). In the right lateral temporal cortex, the mean Kendall $\tau$ was 0.319±0.022 in the group with hippocampal amnesia compared to 0.370±0.020 in the control group. The between-group difference was not significant ($t_{(34)}$ = 1.770, $p$ = 0.086, Cohen's d = 0.590). These results indicate that there were no statistically significant differences between the hippocampal amnesia and control groups in either the left or right LTC, with small to moderate effect sizes observed.

**Ventromedial prefrontal cortex (vmPFC)**

In the left vmPFC, the mean Kendall $\tau$ = 0.284±0.026 in the group with hippocampal amnesia and was 0.306±0.016 in the control group, which was not significantly different ($t_{(34)}$ = 0.714, $p$ = 0.480, Cohen's d = 0.238). In the right vmPFC, the mean Kendall $\tau$ =0.269±0.024 in the group with hippocampal amnesia and was 0.284±0.017 in the control group. Similarly, no significant difference was found between the groups in the right vmPFC ($t_{(34)}$ = 0.522, $p$ = 0.605, Cohen's d = 0.174). These results indicate that there were no statistically significant differences between the hippocampal amnesia and control groups in either the left or right vmPFC, with small to moderate effect sizes observed.

**Parahippocampal cortex (PHC)**

In the left parahippocampal cortex, the mean Kendall $\tau$ = 0.284±0.026 in the group with hippocampal amnesia, whereas it was 0.306±0.016 in the control group, and the between-group difference was not significant ($t_{(34)}$ = 0.714, $p$ = 0.480, Cohen's d = 0.238). In the right PHC, the mean Kendall $\tau$=0.243±0.021 and 0.254±0.017 in the hippocampal amnesia and control groups, respectively, that was no significantly different between-groups ($t_{(34)}$ = 0.415, $p$ = 0.680, Cohen's d = 0.138). These results indicate that there were no statistically significant differences between the hippocampal amnesia and control groups in either the left or right PHC, with small to moderate effect sizes observed.

### 2. Model fit: representation of specific episodic memories

#### Anterior medial prefrontal cortex

#### Left anterior medial prefrontal cortex (amPFC)

In the left amPFC, the mean Kendall τ = 0.0266±0.004 and 0.0295±0.005 in the group with hippocampal amnesia and control groups, respectively. No significant difference was found between the groups ($t_{(34)}$ = 0.447, $p$ = 0.657, Cohen's d = 0.149). In the right amPFC, the mean Kendall τ = 0.0210±0.015 and 0.0308±0.022 in the group with hippocampal amnesia and control groups, respectively. The between-group difference did not reach significance ($t_{(34)}$ = 1.549, $p$ = 0.131, Cohen's d = 0.516). These results indicate that there were no statistically significant differences between the hippocampal amnesia and control groups in either the left or right amPFC, with small to moderate effect sizes observed.

#### Posterior cingulate cortex (PCC)

For the left PCC, the mean Kendall τ = 0.0519±0.005 and 0.0561±0.047 in the group with hippocampal amnesia and control groups, respectively. No significant difference was found between the groups ($t_{(34)}$ = -0.607, $p$ = 0.548, Cohen's d = -0.202). In the right PCC, the mean Kendall τ was 0.0496±0.004 and 0.0523±0.005 in the hippocampal amnesia and control groups. Once again, the between-group difference was not significant ($t_{(34)}$ = 0.503, $p$ = 0.618,Cohen's d = 0.168). These results indicate that there were no statistically significant differences between the hippocampal amnesia and control groups in either the left or right PCC, with small to moderate effect sizes observed.

#### Temporoparietal junction (TpJ)

In the left TpJ, the mean Kendall τ = 0.0434±0.005 and 0.0490±0.006 in the group with hippocampal amnesia and control groups, respectively. The between-group difference was not significant ($t_{(34)}$ = 0.680, $p$ = 0.501, Cohen's d = 0.227). In the right TpJ, the mean Kendall τ was 0.0424±0.005 and 0.0472±0.006 for the amnesic and control groups, respectively. There was no significant difference between-groups ($t_{(34)}$ = 0.611, $p$ = 0.545, Cohen's d = 0.204). These findings suggest minimal differences in TpJ function between group with hippocampal amnesia and control groups, although the results for the left hemisphere should be interpreted with caution due to the lack of a significant model fit in the hippocampal amnesia group.

#### Angular gyrus

In the left angular gyrus, the mean Kendall τ's = 0.0544±0.004 and 0.0647±0.004 in the hippocampal amnesia and control groups, respectively, and was not significantly different between groups ($t_{(34)}$ = 1.868, $p$ = 0.070, Cohen's d = 0.623).

**Left orbitofrontal cortex (OFC)**

In the left OFC, the mean Kendall τ's = 0.0266±0.004 and 0.0327±0.004 in the group with hippocampal amnesia and control groups, respectively, and the between-group difference was not significant ($t_{(34)}$= 1.126; $p$ = 0.268, Cohen's d = 0.375). These results indicate that there were no statistically significant differences between the hippocampal amnesia and control groups in either the left OFC, with small to moderate effect sizes observed.

**Precuneus**

In the left precuneus, the mean Kendall τ's = 0.0559±0.005 and 0.0588±0.004 in the hippocampal amnesia and control groups, respectively, the between-group difference was not significant. ($t_{(34)}$ = 0.437, $p$ = 0.665, Cohen's d = 0.146). In the right precuneus, the mean Kendall τ's was 0.0523±0.005 and 0.0582±0.005 in the hippocampal amnesia and control groups, respectively. No significant between-group difference was found ($t_{(34)}$ = 0.834, $p$ = 0.410, Cohen's d = 0.278). These results indicate that there were no statistically significant differences between the hippocampal amnesia and control groups in either the left or right precuneus, with small to moderate effect sizes observed.

**Retrosplenial cortex**

In the left retrosplenial cortex, the group of participants with hippocampal amnesia exhibited a mean Kendall τ = 0.0218±0.004, whereas control participants exhibited a mean Kendall τ = 0.0224±0.004. No significant difference was found between groups ($t_{(34)}$ = 0.113, $p$ = 0.911, Cohen's d = 0.038). In the right retrosplenial cortex, the mean Kendall τ's=0.0244±0.005 and 0.0235±0.005 in the hippocampal amnesia and control groups, respectively. Similarly, no statistically significant difference was found ($t_{(34)}$ = -0.135, $p$ = 0.894, Cohen's d = -0.045). These results indicate that there were no statistically significant differences between the hippocampal amnesia and control groups in either the left or right retrosplenial cortex, with small to moderate effect sizes observed.

**Middle frontal gyrus (MFG)**

In the left MFG, the mean Kendall τ's = 0.0420±0.004 and 0.0515±0.004 in the hippocampal amnesia and control groups, respectively, whereas the between group difference was not significant ($t_{(34)}$ = 1.768, $p$ = 0.086, Cohen's d = 0.589). In the right MFG, the mean Kendall τ's=0.0371±0.005 and 0.0461±0.004 in the hippocampal amnesia and control group, respectively ($t_{(34)}$ = 1.371, $p$ = 0.179, Cohen's d = 0.457). These results indicate that there were no statistically significant differences between the hippocampal amnesia and control groups in either the left or right MFG, with small to moderate effect sizes observed.

**Lateral temporal cortex (LTC)**

In the left LTC, the mean Kendall τ's = 0.0400±0.005 and 0.0449±0.006 in the hippocampal amnesia and control groups, respectively. The between group difference was not significant ($t_{(34)}$ = 0.636, $p$ = 0.529, Cohen's d = 0.212). In the right LTC, the mean Kendall τ's = 0.0225±0.005 and 0.0463±0.005 in the hippocampal amnesia and the control groups, respectively. The between-group was not significant ($t_{(34)}$ = 1.224, $p$ = 0.230, Cohen's d = 0.408). These results indicate that there were no statistically significant differences between the hippocampal amnesia and control groups in either the left or right LTC, with small to moderate effect sizes observed.

**Ventromedial prefrontal cortex (vmPFC)**

In the left vmPFC, the mean Kendall τ's = 0.0271±0.004 and 0.0315±0.005 in the hippocampal amnesia and control groups, respectively. No significant difference between groups was found ($t_{(34)}$ = 0.709, $p$ = 0.484, Cohen's d = 0.236). In the right vmPFC, the mean Kendall τ's = 0.0233±0.004 and 0.0285±0.005 in the hippocampal amnesia and the control groups respectively, with no significant difference between the groups ($t_{(34)}$ = 0.710, $p$ = 0.482, Cohen's d = 0.237). These results indicate that there were no statistically significant differences between the hippocampal amnesia and control groups in either the left or right vmPFC, with small to moderate effect sizes observed.

**Parahippocampal cortex (PHC)**

In the left PHC, the mean Kendall τ's = 0.0223±0.004 and 0.0171±0.005 in the hippocampal amnesia and control groups, respectively, with no significant difference between-groups ($t_{(34)}$ = -0.861, $p$ = 0.395, Cohen's d = -0.287). In the right PHC, the mean Kendall τ's = 0.0196±0.004 and 0.0150±0.004 in the hippocampal amnesia and the control groups, respectively. No significant difference was found ($t_{(34)}$ = -0.804, $p$ = 0.427, Cohen's d = -0.268). These results indicate that there were no statistically significant differences between the hippocampal amnesia and control groups in either the left or right PHC, with small to moderate effect sizes observed.
